# High-resolution single-cell atlas reveals diversity and plasticity of tissue-resident neutrophils in non-small cell lung cancer

**DOI:** 10.1101/2022.05.09.491204

**Authors:** Stefan Salcher, Gregor Sturm, Lena Horvath, Gerold Untergasser, Georgios Fotakis, Elisa Panizzolo, Agnieszka Martowicz, Georg Pall, Gabriele Gamerith, Martina Sykora, Florian Augustin, Katja Schmitz, Francesca Finotello, Dietmar Rieder, Sieghart Sopper, Dominik Wolf, Andreas Pircher, Zlatko Trajanoski

**Author notes:** Correspondence (A.P.), (Z.T.). These authors contributed equally.

## Abstract

Non-small cell lung cancer (NSCLC) is characterized by molecular heterogeneity with diverse immune cell infiltration patterns, which has been linked to both, therapy sensitivity and resistance. However, full understanding of how immune cell phenotypes vary across different patient and tumor subgroups is lacking. Here, we dissect the NSCLC tumor microenvironment at high resolution by integrating 1,212,463 single-cells from 538 samples and 309 patients across 29 datasets, including our own dataset capturing cells with low mRNA content. Based on the cellular composition we stratified patients into immune deserted, B cell, T cell, and myeloid cell subtypes. Using bulk samples with genomic and clinical information, we identified specific cellular components associated with tumor histology and genotypes. Analysis of cells with low mRNA content uncovered distinct subpopulations of tissue-resident neutrophils (TRNs) that acquire new functional properties in the tissue microenvironment, providing evidence for the plasticity of TRNs. TRN-derived gene signature was associated with anti-PD-L1 treatment failure in a large NSCLC cohort.

**In brief:** Salcher, Sturm, Horvath et al. integrate single-cell datasets to generate the largest transcriptome atlas in NSCLC, refining patient stratification based on tumor immune phenotypes, and revealing associations of histological subtypes and genotypes with specific cellular composition patterns.

Coverage of cells with low mRNA content by single-cell sequencing identifies distinct tissue-resident neutrophil subpopulations, which acquire new properties within the tumor microenvironment. Gene signature from tissue-resident neutrophils is associated with immune checkpoint inhibitor treatment failure. The integrated atlas is publicly available online (https://luca.icbi.at), allowing the dissection of tumor-immune cell interactions in NSCLC.

**Highlights:**

- High-resolution single-cell atlas of the tumor microenvironment (TME) in NSCLC.
- Histological tumor subtypes and driver genes imprint specific cellular TME patterns.
- scRNA-seq of cells with low transcript count identifies distinct tissue-resident neutrophil (TRN) subpopulations and non-canonical functional properties in the TME niche.
- TRN gene signature identifies patients who are refractory to treatment with PD-L1 inhibitors.

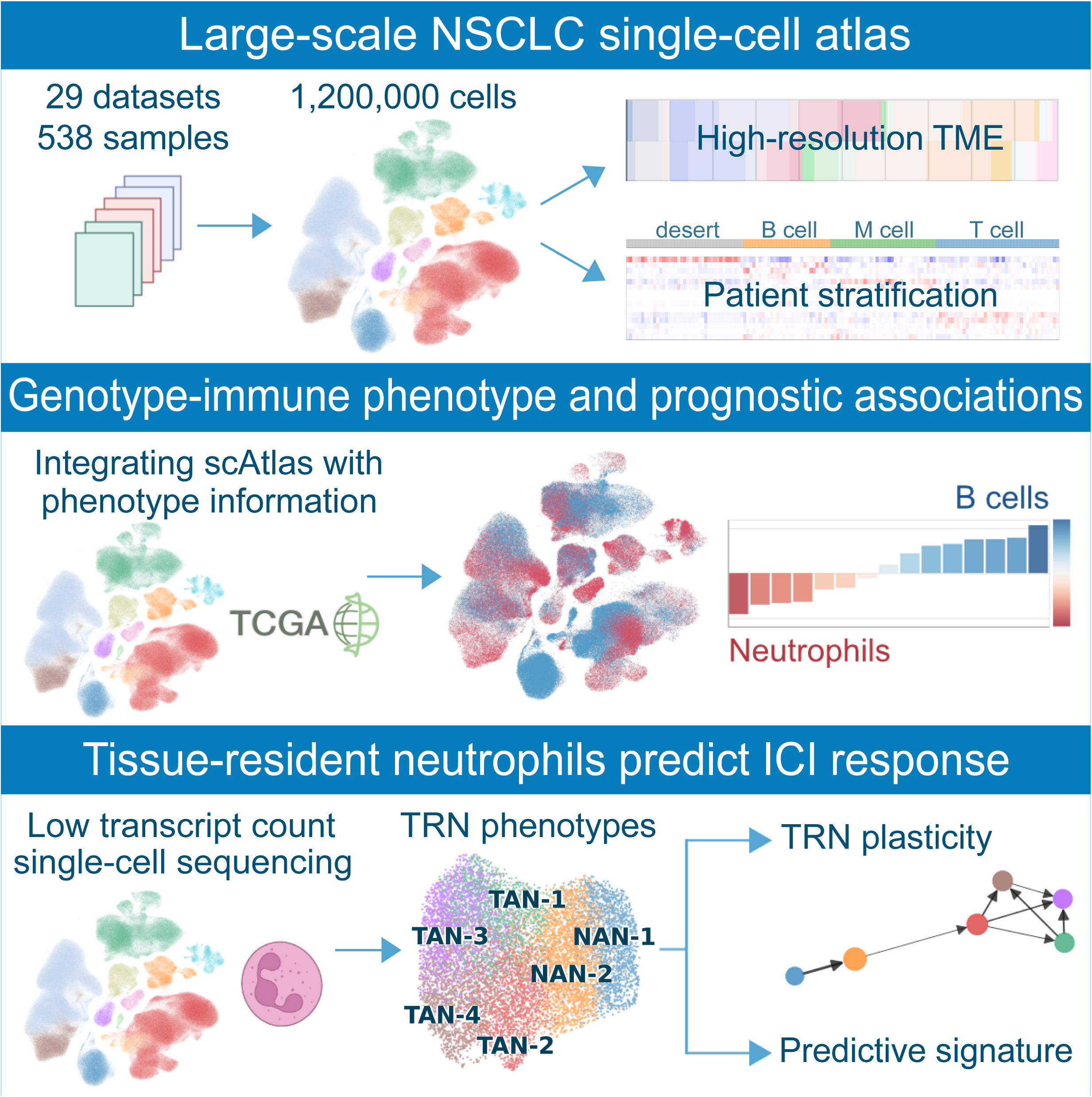

## INTRODUCTION

Single-cell RNA sequencing (scRNA-seq) allows transcriptome dissection at single-cell resolution and has revolutionized our understanding of diverse host microenvironments in health and disease (Potter, 2018). Recent technological and bioinformatic advances have made this technology increasingly accessible to scientists, providing highly anticipated applications ranging from basic science to clinical biomarker discovery (Potter, 2018) and identification of drug resistance mechanisms (Aissa et al., 2021). Non-small cell lung cancer (NSCLC) is a highly aggressive and heterogenous disease with diverse histological subtypes and distinct mutational signatures (Chen et al., 2014), which accounts for an annual global death rate of 1.8 Mio patients (Sung et al., 2021). The technical advances in scRNA-seq technologies enabled the dissection of the complex NSCLC tumor microenvironment (TME) in different stages and numerous scRNA-seq NSCLC studies have identified a hitherto underestimated TME heterogeneity in early and advanced disease (Chen et al., 2020; Goveia et al., 2020; Kim et al., 2020; Lambrechts et al., 2018; Laughney et al., 2020; Leader et al., 2021; Maier et al., 2020; Maynard et al., 2020; Wu et al., 2021; Xing et al., 2021; Zilionis et al., 2019). Furthermore, these studies highlighted the importance of small cell populations in governing essential biological pathways such as immune cell activation or trafficking by tumor endothelial cells (Goveia et al., 2020), and the key immunomodulatory role of fibroblasts (Lambrechts et al., 2018). These studies underpinned the multifactorial disease character of NSCLC, with both inter- as well as intra-individual heterogeneities. However, a major limitation of these studies is the limited number of analyzed patient samples per study. Moreover, the lack of genomic data as well as long-term follow-up data prevents comprehensive dissection of the biological heterogeneity and its potential contribution to therapy resistance and survival outcome.

While scRNA-seq helped to provide a considerable depth in mapping the composition of the NSCLC TME, technical and methodological variations between the different studies (e.g. interlaboratory differences in cell handling, scRNA-seq platforms, sequencing techniques, data-processing algorithms) result in significant inconsistencies and knowledge gaps. As such, not all textbook-defined cell types (e.g. neutrophilic granulocytes) have been portrayed in the same depth and extension yet, posing an unmet need to characterize these populations as well. In NSCLC, it is well accepted that next to cancer cells, leukocytes compose the majority of cells within the TME (Stankovic et al., 2018; Wu et al., 2021). Particularly since immunotherapy is routinely used in clinical practice, in-depth characterization of the cancer immune cell compartment has been intensively pushed forward and diverse cellular subsets have been profiled (Chen et al., 2020; Guo et al., 2018; Maier et al., 2020). Previous compositional analyses by flow cytometry as well as histological work-up suggested that neutrophils compose a significant proportion of all tumor-resident leukocytes, with an estimated abundance ranging from 8 to 20% (Eruslanov et al., 2014; Kargl et al., 2017; Stankovic et al., 2018). Intriguingly, when looking at the numerous scRNA-seq studies in NSCLC published over the last years (Chen et al., 2020; Goveia et al., 2020; Guo et al., 2018; He et al., 2021; Kim et al., 2020; Lambrechts et al., 2018; Laughney et al., 2020; Maynard et al., 2020; Wu et al., 2021; Xing et al., 2021; Zilionis et al., 2019), neutrophils are clearly underrepresented. This discrepancy is most likely based on technical issues rather than on biological phenomena, but its clarification is of immense importance for both, our fundamental immunological understanding of NSCLC and for potential translational clinical investigations. This notion is underscored by pre-clinical data suggesting that neutrophils are essential mediators of both pro- and anti-tumor inflammatory pathways (reviewed in (Shaul and Fridlender, 2019)) as well as by correlative studies in NSCLC patients linking the neutrophil/lymphocyte ratio with clinical outcome and response to immunotherapy (Kargl et al., 2019; Peng et al., 2015; Templeton et al., 2014).

To overcome the above-mentioned hurdles, we here present a comprehensive NSCLC scRNA-seq atlas that integrates major publicly available datasets as well as a dataset we have generated that includes cells with very low transcript count, resulting in a large data resource covering 223 NSCLC patients and 86 healthy controls. Using this atlas and cutting-edge computational tools, we portray the TME in NSCLC at single-cell resolution and stratify the patients into distinct immune phenotypes. Furthermore, integration of bulk RNA-seq data showed that histological tumor subtypes and driver genes imprint specific cellular TME patterns. Deep characterization of tissue-resident neutrophils (TRNs) uncovered distinct subpopulations of tumor-associated neutrophils (TANs) and normal adjacent tissue-associated neutrophils (NANs), and showed that TRN acquire non-canonical functional properties in the TME niche. Finally, we derived a TRN gene signature that associates with anti-PD-L1 treatment failure in a large NSCLC cohort.

## RESULTS

### Generation of a core large-scale NSCLC single-cell atlas

We first developed a core NSCLC atlas by compiling scRNA-seq data from 19 studies and 21 datasets comprising 505 samples from 298 patients (Figure 1A). This comprehensive NSCLC single-cell atlas integrates expert-curated, quality-assured and pre-analyzed transcriptomic data from publicly available studies as well as our own dataset (UKIM-V) in early and advanced stage NSCLC of any histology (Chen et al., 2020; Goveia et al., 2020; Guo et al., 2018; He et al., 2021; Kim et al., 2020; Lambrechts et al., 2018; Laughney et al., 2020; Maynard et al., 2020; Mayr et al., 2021; Wu et al., 2021; Zilionis et al., 2019) and 7 studies for control purpose (Adams et al., 2020; Habermann et al., 2020; He et al., 2021; Madissoon et al., 2019; Reyfman et al., 2019; Travaglini et al., 2020; Vieira Braga et al., 2019). The selected studies were published between July 2018 and May 2021 and the incorporated datasets were generated with six different sequencing platforms, including the most commonly applied 10x Chromium (10x Genomics) as well as Smart-seq2 (Picelli et al., 2013), GEXSCOPE (Singleron), inDrop (Klein et al., 2015) and Drop-Seq (Macosko et al., 2015). We further integrated our own data generated with the microwell-based BD Rhapsody scRNA-seq platform (BD Biosciences). We specifically selected studies using comparable protocols for sample processing and data generation, such as sequencing of whole cells. We did not exclude studies that applied flow cytometry-based cell-sorting prior to sequencing, as these incorporate relevant information on rare cell types (Goveia et al., 2020; Guo et al., 2018; Maier et al., 2020). From non-NSCLC studies we exclusively included those parts of the published data that were relevant for our atlas: from the Maddisoon dataset (Madissoon et al., 2019) we only included lung samples, from Adams (Adams et al., 2020), Reyfman (Reyfman et al., 2019), Haberman (Habermann et al., 2020), Vieira Braga (Vieira Braga et al., 2019), and Mayr (Mayr et al., 2021) datasets we only used the control samples (including normal lung tissue of tumor patients, which we termed “normal_adjacent” or lung tissue of organ donors without history of pulmonary disease). From the Adams et al. dataset we also included data from patients (n=18) with chronic obstructive pulmonary disease (COPD) as chronic inflammatory pulmonary disease cohort with an increased lung cancer risk. Important study characteristics are summarized in Table S1.

**Figure 1.**
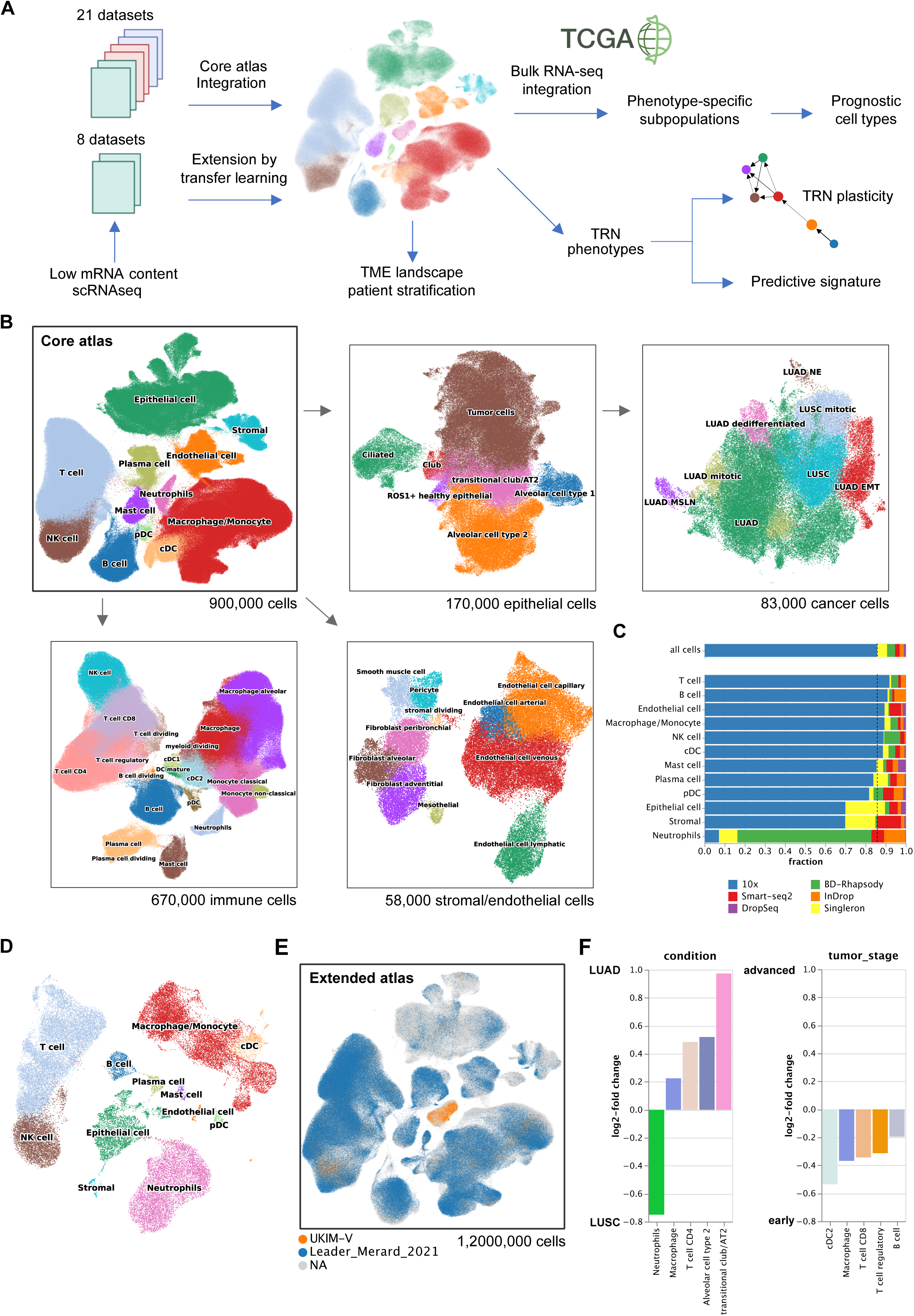
Schematic outline of the overall concept used in this study. (A) Summary of the data integration and analysis workflow. (B) Overview of the core NSCLC atlas and the epithelial, immune, and stromal/endothelial components depicted as UMAP plots. (C) Fractions of depicted cell types per scRNA-sequencing platform. (D) UMAP of the UKIM-V dataset (n=8) colored by cell-types. (E) Core atlas extended by Leader (Leader et al., 2021) and UKIM-V-2 datasets. (F) Cell type composition by tumor type (LUAD, LUSC) and stage (early, late) calculated with scCODA (Bayesian model for differential composition analysis) using tumor cells as reference cell-type (assumed to be constant between conditions) and 500,000 Markov-chain monte carlo iterations. FDR=0.1.

In total, the core atlas includes transcriptomic data from 212 NSCLC patients and 86 control individuals, comprising 196 tumor samples and 168 non-tumor control samples. Of the 212 NSCLC patients, 156 were clinically annotated as lung adenocarcinoma (LUAD), 41 as lung squamous cell carcinoma (LUSC), and 15 were not otherwise specified (NSCLC NOS). NSCLC samples include tissue of the primary tumor or metastasis (adrenal, brain, lymph node, liver, pleura or pleural effusion) that were obtained either by surgical resection (most cases) or by computed tomography (CT)- and bronchoscopy-guided biopsies. We clustered the NSCLC patients’ disease stages as early (UICC stage I – II) *versus* advanced (UICC III-IV) disease, as not all studies provided sufficient information on tumor stages. Among the control samples, 89 were derived from distant non-malignant tissue of lung tumor patients (annotated as normal_adjacent), of which 65 have a patient-matched tumor sample. Further 10 samples were derived from non-tumor affected lymph nodes of NSCLC patients (annotated as normal), and 79 samples from patients without evident lung cancer history (annotated as normal). Of the control patients 18 had a history of COPD. Overall, the atlas integrates 898,422 single cells, which are annotated to 12 coarse cell type identities and 38 major cell subtypes or cell states (e.g. dividing cells) based on previously established canonical single-cell signatures (Figure S1A) including 169,223 epithelial cells (alveolar cells type 1 and 2, club cells, transitional club/AT2, ciliated cells, ROS1^+^ healthy epithelial, and tumor cells), 670,409 immune cells (T cells, B cells, dendritic cells (DC) including the subtypes mature, conventional and plasmacytoid DCs, mast cells, macrophages, monocytes, neutrophils, plasma cells, natural killer (NK) cells), and 58,790 stromal cells (fibroblasts (adventitial, peribronchial, alveolar) and endothelial cells (lymphatic, capillary, venous, arterial), pericytes, smooth muscle cells, mesothelial cells) (Figure 1B). After harmonization of cell-type clusters between datasets (see methods), we found high consistency of cell type-specific markers across the different scRNA-seq datasets and patients (Figure S1B). For some cell types, the relative composition varied markedly among the different datasets (e.g. stromal cells, epithelial cells, neutrophils) (Figure S1B). Overall, tumor and control samples showed different cell-type composition patterns (Figure S1C). The cell type composition for each data set, the tissue of origin, and the patients within the core atlas are shown in Figure S1B.

Tumor classification relies on morphology (small cell, large cell) and differentiation status (adeno=LUAD, squamous=LUSC). However, NSCLC tumors comprise very heterogenous cellular composition with mixed characteristics. Previous scRNA-seq studies have categorized the major and clinically relevant types of LUSC and LUAD. To the best of our knowledge, no transcriptomic signatures have been published yet that would allow for a more precise subclassification. Malignant epithelial cells (in total 83,439 cells) in the atlas showed high heterogeneity of the transcriptomic profiles. Due to the large patient cohort we were able to apply a comprehensive lung tumor cell classification based on their specific marker gene expression signatures (Figure S1D), depicting LUSC (*KRT5, KRT6A, KRT17, SOX2, NTRK2, TP63*) and LUAD (*CD24, MUC1, KRT7, NAPSA, NKX2-1, MSLN*) as well as LUAD with signs of epithelial-to-mesenchymal transition (LUAD EMT) (*VIM, SERPINE1, CDH1, MIF*), LUAD with neuroendocrine features (LUAD NE) (*CHGA, SYP, NCAM1, TUBA1A*), LUAD with high expression of mesothelin (*MSLN*) associated with EMT and metastasis (LUAD MSLN) (He et al., 2017), and dedifferentiated carcinoma expressing both, LUAD and LUSC markers (*TACSTD2, ARG2*) (Figure 1B). There are highly mitotic/proliferative clusters (*TOP2A, MKI67*) of both LUAD and LUSC that may resemble highly aggressive and invasive tumor cells. The LUAD EMT cells likely resemble an invasive, pro-metastatic cluster characterized by the plasminogen activator PAI1 (*SERPINE1*) or the mesenchymal protein vimentin (*VIM*) that are both involved in cell adhesion, invasion and angiogenesis (Francart et al., 2020; Kubala et al., 2018). Besides, all subclusters showed a high expression of the conserved non-coding RNAs *MALAT1* and *NEAT1* (Figure S1D) that were previously linked to metastasis formation in NSCLC (Ji et al., 2003; Qi et al., 2018).

### Neutrophils are underrepresented in most scRNA-seq studies

In both tumor and normal lung tissue of all core atlas samples, the neutrophil cluster (*FCGR3B, CSF3R, CXCR2* and *G0S2*) comprised 8,468 cells with overt low mRNA counts. Overall, neutrophils account for only 1% percent of all atlas cells (Figure S1E). Remarkably, more than 60% of all atlas neutrophils derive from the UKIM-V dataset (Figure 1C, Figure S1B), in which neutrophils compose 21% of all cells and 24% of all leucocytes, respectively. The remaining neutrophil data originate mainly from four other datasets (Chen et al., 2020; Maynard et al., 2020; Wu et al., 2021; Zilionis et al., 2019), in all of which neutrophils comprise less than 6% of all leucocytes. Particularly in those studies using the droplet-based 10x Chromium platform, neutrophils are completely absent or only rarely depicted (Figure S1F). Our comparative flow cytometry analysis demonstrated that neutrophils account for 10-20% of all leucocytes in NSCLC tumor and normal adjacent tissues (n= 63) (Figure S1G), which is in accordance with previous flow cytometry and histology data (Eruslanov et al., 2014; Kargl et al., 2017; Stankovic et al., 2018). Thus, low neutrophil abundance seen in previous scRNA-seq datasets suggests an underrepresentation, most likely due to technical issues. Neutrophils are fragile, short-lived cells (circulatory half-life of 7-10 hours in humans (Shaul and Fridlender, 2019)) and particularly sensitive to handling procedures. In addition, neutrophils express an exceptionally low amount of mRNA molecules (Zilionis et al., 2019), which impedes their recovery in scRNA-seq data.

Comparative analysis of scRNA-seq platforms indicated that the BD Rhapsody workflow captures a notably high number of mRNA molecules per cell and may thus be especially suitable to depict low-mRNA content cells (Figure S1H). Concordantly, in-house benchmarking of high-throughput scRNA-seq platforms support the notion that the BD Rhapsody detects relatively high numbers of mRNA molecules per cell thus enhancing the recovery of low-mRNA-content cells (data not shown). As a consequence, neutrophils represented a major cell cluster in the UKIM-V dataset generated with BD Rhapsody, whereas the low-mRNA content neutrophil cluster could not be detected in the datasets generated with 10x Chromium, and only to a very limited extend in the GEXSCOPE/Singleron, InDrop, DropSeq and Smart-seq2 datasets (Figure 1C). These divergent results may be attributed to technical differences between platforms, including the different mRNA count numbers but also applied cell handling strategies, such as FACS-sorting that may *per se* diminish neutrophils due to their inherent fragility (as done in the Smart-seq2 protocol).

### scRNA-seq of low-mRNA content cells

Due to high clinical relevance of neutrophils and the need for their better in-depth transcriptomic characterization, we used the advantage of the BD Rhapsody platform in depicting cells with low mRNA content for further analysis. As our initial dataset included only three patients, we next performed scRNA-seq of additional five NSCLC patients to increase the statistical weight of our cohort (UKIM-V cohort). In total our dataset contains tumor and adjacent normal lung tissue from eight patients (3x male, 5x female) undergoing lobectomy for early-stage treatment-naïve NSCLC (5x LUAD, 3x LUSC). Cells were freshly isolated, processed, and sequenced as described in detail in the methods section. The UKIM-V dataset (1 and 2) comprises 47,485 cells that cluster into all main lung cell types defined by the expression of specific marker genes (Figure 1D). Neutrophils are characterized as cell cluster with exceptionally low mRNA content and thus a relatively low number of detected transcripts, but due to the relatively high number of mRNA molecules captured per cell (UMI counts in epithelial cells: 10,433) we could readily depict these cells in the UKIM-V dataset. The 8,313 identified neutrophils identified in the UKIM-V dataset were derived from control lung (n=2,677) and corresponding tumor tissue (n=5,636).

### Extension of the core single-cell atlas by transfer learning

Recently developed transfer-learning method scArches (Lotfollahi et al., 2022) enables the extension of the core atlas using additional existing datasets as well as datasets from future studies. We therefore mapped one recently published dataset comprising 288,157 cells (Leader et al., 2021) as well as our second UKIM-V dataset onto the atlas (Figure 1E). The cell type annotations were transferred from the core atlas to the two new datasets on the basis of transcriptomic similarity in the batch-corrected joint embedding (see Methods). The resulting extended atlas integrates 29 datasets from 19 studies and comprises 1,212,463 cells, 38 cell types, 538 samples, and 309 patients, resulting in 1.62 billion expression values. The overall cell type composition of the extended atlas is shown in Figure S1I. All subsequent analysis were carried out on the extended atlas data set unless otherwise specified.

Identification of changes in cell type compositions in distinct histological or genetic tumor types and tumor stages is of utmost importance as it can highlight hetero-cellular interactions and possibly enables association(s) with therapy response. However, detecting shifts in cell type composition using scRNA-seq data is challenging due to the inherent bias present in cell type compositions and low sample sizes. We therefore adopted a Bayesian model for identifying changes in cell type compositions while controlling for the false discovery rate (scCODA tool (Buttner et al., 2021)). scCODA requires setting a reference cell type that is assumed to be constant between conditions. When comparing cellular composition in LUSC and LUAD with tumor cells as reference cell type, we found a significant higher proportion of neutrophils in LUSC, whereas macrophages, alveolar cells type 2, CD4^+^ T cells and transitional club/AT2 were more abundant in LUAD (Figure 1F), underlining histology-dependent differences in TME characteristics. The analysis of the cell type compositions in early stage compared to advanced stage NSCLC tumors showed higher abundance of immune cells (CD8^+^, CD4^+^, T regulatory cells, and cDC2) (Figure 1F).

### Single-cell composition of the TME reveals distinct NSCLC tumor immune phenotypes

Next, we stratified NSCLC patients based on infiltration patterns of their TME using the extended atlas. Unsupervised clustering on batch-corrected cell-type fractions revealed four distinct tumor immune phenotypes (Figure 2A): 1) immune deserted tumors (ID; i.e. no significant immune cell infiltration but high tumor cell fraction); 2) the subtype of tumors with B cell dominance (B; B cell, plasma cell, mast cell); 3) the subtype of tumors with myeloid dominance (M; macrophage/monocyte) infiltration; and 4) subtype of tumors with T cell dominance (T; CD8^+^, CD4^+^, T regulatory cells). The affiliation of UKIM-V patients to myeloid cell and T cell subtypes was validated using flow cytometry and histology (Figure S2A). Across the strata, most patients of the B cell, myeloid cell, and T cell subtypes showed LUAD histology, whilst half of the LUSC patients were over-represented in the immune deserted subtype (Figure S2B). Neutrophils were excluded from patient stratification since they are underrepresented in most datasets. However, previous studies showed a relative enrichment of neutrophils and M2 macrophages in otherwise immune deserted tumors (Park et al., 2022). Using logistic regression, we did not find any association of the different patient strata to tumor stages (early *vs.* advanced) or sex.

**Figure 2.**
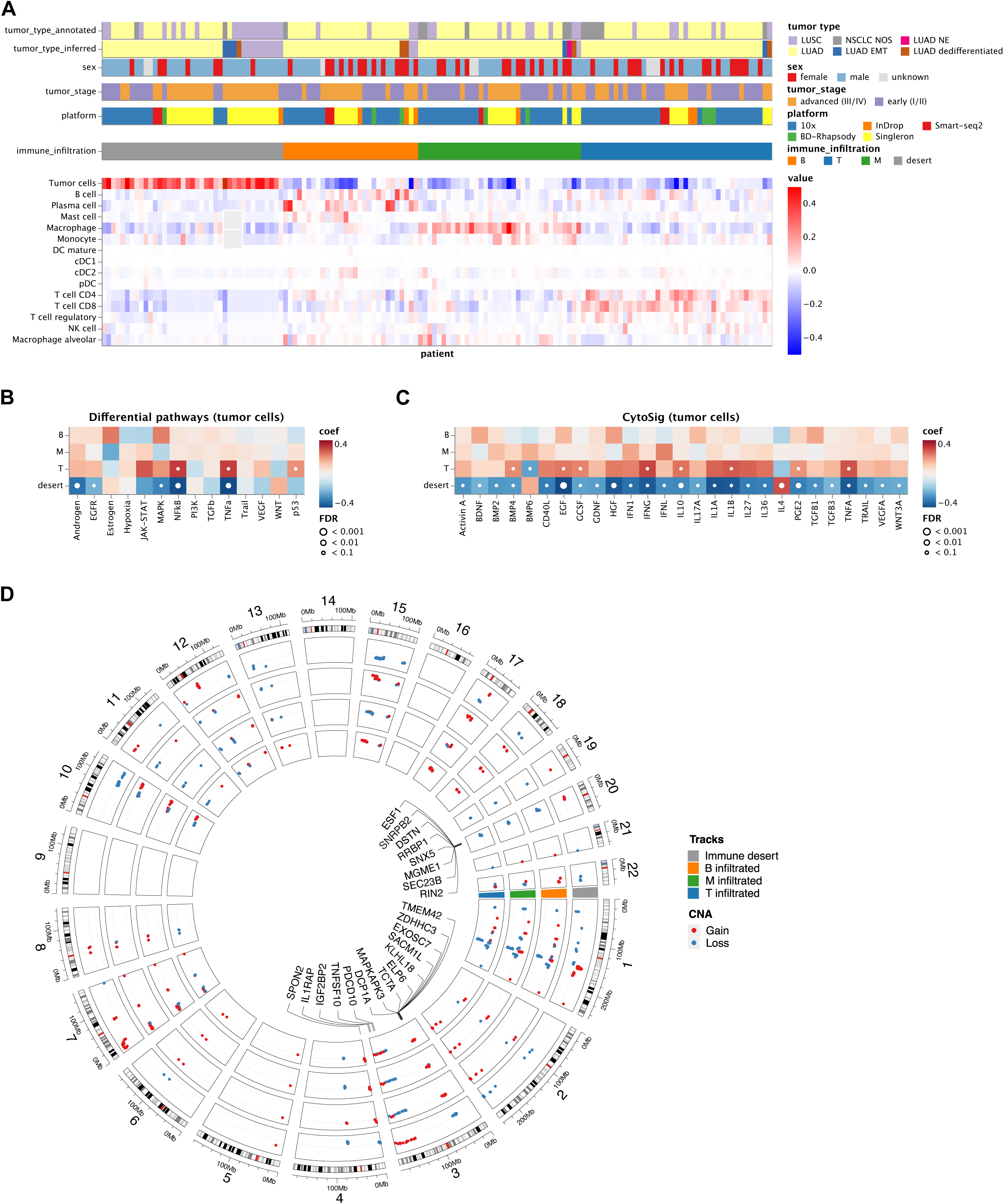
Tumor immune phenotypes in NCSCL. (A) Patient characteristics and stratification of the tumor immune phenotypes. *tumor_type_annotated* refers to the histological subtypes as provided by the original datasets based on pathological assessment; *tumor_type_inferred* is based on the most abundant tumor cell-type in the scRNA-seq atlas. Immune phenotypes are derived by unsupervised, graph-based clustering of batch-corrected cell-type fractions with Pearson correlation as distance metric. (B+C) Differential activation of (B) PROGENy cancer pathways-(C) CytoSig cytokine signalling signatures in tumor cells between the four tumor immune phenotypes. Heatmap colors indicate the deviation from the overall mean, independent of tumor histology and stage. White dots indicate significant interactions at different false-discovery-rate (FDR) thresholds. P-values have been calculated using a linear model f-test. Only cytokine signatures with an FDR < 0.1 in at least one patient group are shown. (D) Circos plot of significant CNAs for the four tumor immune phenotypes of the single-cell NSCLC cohort. Highlighted are selected genes that matched the CNAs in the TCGA cohort. Significant gains/losses (FDR<0.1) were determined using a linear model with HC3 standard errors.

To identify tumor-cell based TME imprinting characteristics, we next analyzed differentially enriched pathways (Schubert et al., 2018) in the tumor cells of each of the four immune phenotypes. The immune deserted subtype showed downregulation (FDR < 0.1) of the immune mediating pathways *NFkB* and *TNFα,* as well as androgen, *EGFR*, and *MAPK* pathways (Figure 2B). Previously, androgen receptor signaling has been shown to suppress PD-L1 transcription in HCC cells and may thus exert immune stimulatory effects (Jiang et al., 2020). Analysis of differentially expressed transcription factors (Garcia-Alonso et al., 2019; Holland et al., 2020) in the tumor cells of each subtype showed a significant down-regulation of *FOXO4* and *PPARA* in the immune deserted subtype (Figure S2C). As previously reported, FOXO transcription factors are essential mediators of immune cell homeostasis (Peng, 2008), whereas activation of peroxisome proliferator activated receptors (PPARs, including PPARA) supports T cell survival in the TME by reprogramming their energy metabolism (Chowdhury et al., 2018). Thus, *FOXO4* and *PPARA* downregulation may promote an immune deserted phenotype.

We then applied the tool *CytoSig* (Jiang et al., 2021) to define enriched cytokine signaling signatures in tumor cells of each immune phenotype (Figure 2C). *CytoSig* analyses defined cytokine signatures that are differentially expressed when a cell is exposed to a specific cytokine (that is name-giving for the respective cytokine signature). Solely the signature of the tolerogenic cytokine IL-4, a modulator of T regulatory cell-mediated immune suppression (Yang et al., 2017), was significantly elevated in the immune deserted subtype (Figure 2C). As expected, we found significant upregulation of most cytokine signatures in the B cell, myeloid cell, and T cell subtypes, including pro-inflammatory (e.g. TNFA, IL1B, CD40L) but also growth factors (EGF, VEGF), underlining immune cells as the major signaling partners in the TME (Figure 2C).

Next, we used the scRNA-seq data to infer copy number alterations (CNAs) in cancer cell populations as well as CNA-based intratumoral heterogeneity scores (ITH_CNA_; see methods). The CNA profiles of the 199 cancer patients showed interas well as and intra-patient heterogeneity. However, detailed analysis of cell subtypes and ITH_CNA_ scores revealed no significant correlation with major immune cell types (data not shown). Moreover, we found no correlation of heterogeneity scores with tumor histology, patient immune infiltration subtype or tumor stage (Figure S2D. To investigate the impact of genetic aberrations on the tumor immune phenotypes we analyzed the gains and losses among the four immune subtypes (Figure 2D). There were 279 genes with significant gains or losses across the four immune subtypes (Table S2). We then annotated patients from the TCGA cohort to the immune deserted and immune infiltrated phenotypes using results from deconvolution analysis from bulk RNA-seq data and analyzed the CNAs. We identified 4984 significant genes with gains or losses across the immune deserted and immune infiltrated phenotypes s (Table S3) of which 56 were detected also in the scRNA-seq data (Table S4). Selected genes from the intersection between the scRNA-seq data and the TCGA data with opposite genetic alterations between the immune deserted subtype and the immune infiltrated subtypes (B cell, T cell, and myeloid cell subgroups) are shown in Figure 2D and S2E. For example, genes on chromosome 3 implicated in cell apoptosis (*PCDC10*, *TNFSF10* (*TRAIL*)) showed gains in the immune deserted subtype and losses in the immune infiltrated groups in both, scRNA cohort and the TCGA cohort.

### Analysis of cell-cell communication reveals hetero-cellular crosstalk in the TME

Using the CellPhoneDB ligand-receptor complexes database, we next assessed differences in the hetero-cellular crosstalk of tumor cells towards diverse immune cells among the two major histotypes LUAD and LUSC, by analyzing the top 15 differentially expressed tumor cell ligands (Figure 3A, top 30 ligands are shown in Figure S3A). Overall, in both histologies tumor cellular interactions were directed to diverse immune cell subtypes with different targets. In LUAD, we found a prominent upregulations of the KDR-VEGFA axis from tumor cells towards neutrophils and mast cells, potentially implicating immunosuppressive signaling by tumor cells in this histotype (Motz and Coukos, 2011). Other major LUAD pathways included ADGRE5-CD55, associated with migration and invasion (Yin et al., 2018), and LGALS9-HAVCR2 (galectin9-TIM3), known to suppress anti-tumor immunity (Goncalves Silva et al., 2017; Yasinska et al., 2019). Conversely in LUSC, pro-migratory signaling involves SPP1 (osteopontin) (Shojaei et al., 2012) that has previously been reported as upregulated in lung cancer tissue particularly in squamous histology (Zhang et al., 2001). Interestingly, neutrophil recruiting signals including ANXA1-FPR1/2 and CCL26-CCR1 (Metzemaekers et al., 2020) are upregulated in LUSC but not LUAD, potentially explaining the higher neutrophil infiltration in LUSC (Figure S5A).

**Figure 3.**
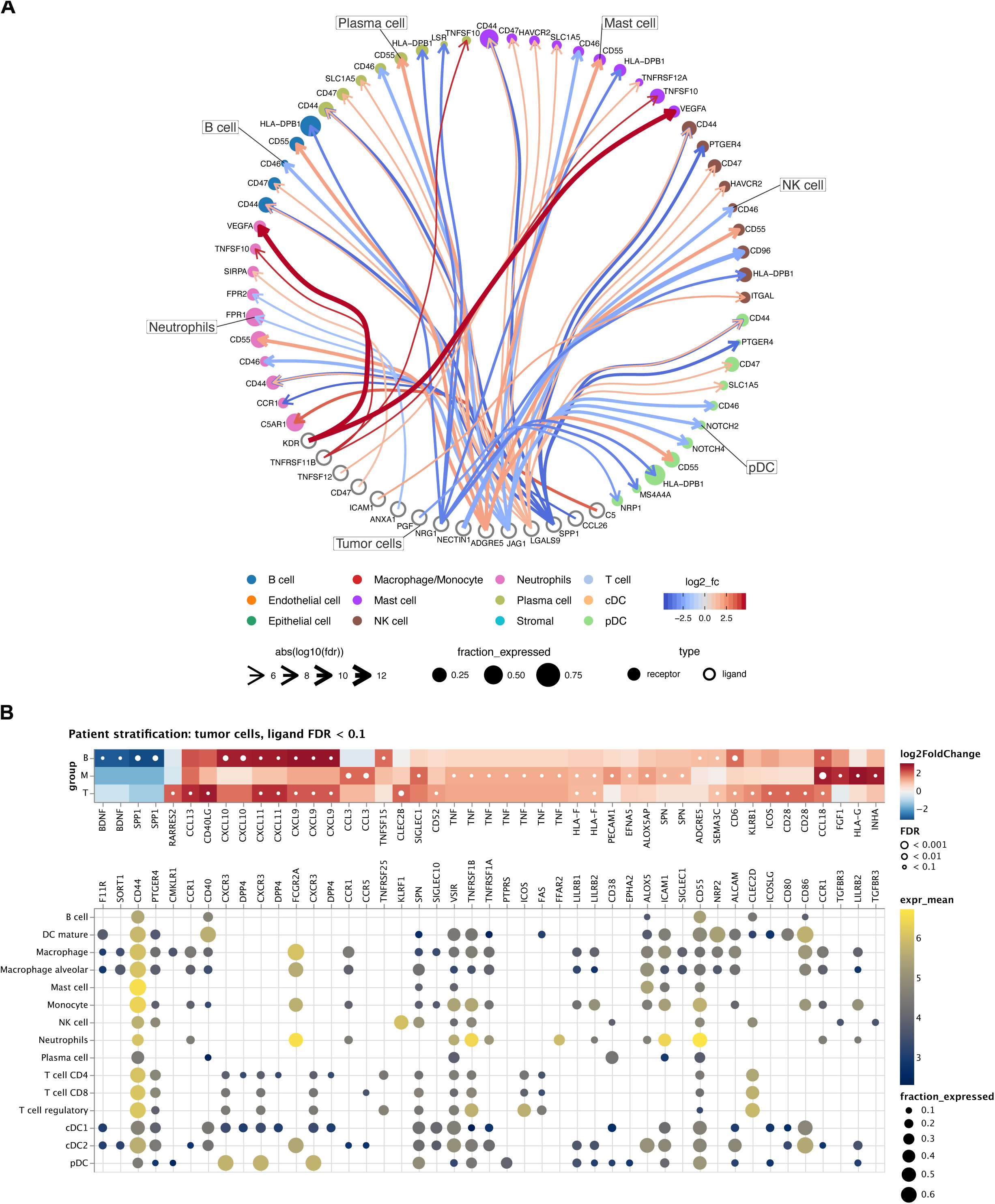
Cellular cross-talk analysis. (A) Circos plot of the cellular crosstalk of tumor cells towards the major immune cells in LUAD versus LUSC.Shown are the top 15 differentially expressed ligands. Red interactions are upregulated in LUAD, blue interactions are upregulated in LUSC. (B) Tumor-immune cell cross-talk in each patient subtype. Upper panel: top 30 differentially expressed ligands of tumor cells in each subtype (B, M, T) compared to immune deserted (DESeq2 on pseudo-bulkFDR<0.1). Bottom panel: Respective receptors and the expression by cell type. Dot sizes and colors refer to the fraction of cells expressing the receptor and gene expression, respectively, averaged over all patients. Dots are only shown for receptors that are expressed in at least 10% of the respective cell-types.

Next, we investigated the crosstalk of tumor cells to immune cells within the patient immune subtypes T cell, B cell, and myeloid cell, and analyzed differentially expressed tumor cell ligands of the immune infiltrated subtypes compared to the immune deserted subtype (Figure 3B). While most significant interactions were found to be upregulated in the immune-infiltrated subtypes, SPP1 signaling was down-regulated when compared to the immune deserted subtype, underlining previous *in silico* analysis that have associated high SSP1 expression with an immune-evasive TME in NSCLC (Zheng et al., 2021). Other significant interactions include upregulated signaling of chemokines CXCL9/10/11 to their cognate receptors DPP4 and CXCR3 in the T and B subtypes, suggesting active chemotaxis by tumor cells to promote immune infiltration (Mikucki et al., 2015). Of note, while T cells are directly involved in this signaling cascade by expression of CXCR3/DDP4, this was not found on B cells, whose involvement in CXCL9-11 is less known.

### Integration of bulk RNA-seq data reveals genotype-immune phenotype associations

scRNA-seq provides an unprecedented view on the cellular heterogeneity in the TME. However, the majority of the scRNA studies lack both, cancer genotype information and survival data. The TCGA reference dataset includes this information together with bulk RNA-sequencing data. Using the recently published computational method SCISSOR (Choi et al., 2021) we evaluated the association of atlas-derived cell type transcriptomic signatures with genotype and survival data from the TCGA reference dataset including 1026 patients (UICC I-IV, LUAD and LUSC). In a previous pan-cancer study using bulk RNAseq data we have shown that genomic features including mutational load, tumor heterogeneity, and specific driver genes determine immune phenotypes (Charoentong et al., 2017). Here, the high-resolution of the single-cell NSCLC atlas enabled for the first-time an in-depth analysis of these determinants. EGFR, TP53, and STK11 mutations showed distinct immune infiltrates (Figure 4A-D, Figure S4A-B). For example, cDC1 and cDC2 showed opposite infiltration patterns in LUAD patients with either EGRF-(high cDC infiltration) or KRAS-mutated tumors (low cDC infiltration) (Figure 4A). Conversely, TP53 and STK11 mutated genotypes were associated with CD8^+^ T infiltration, which is not seen in EGFR-or KRAS-mutated tumors (Figure 4C-D, Figure S5A-B). Interestingly, KRAS mutations were associated with stromal and endothelial cell infiltration that is not seen in the other genotypes examined. Also, in tumors with EGFR, TP53 and STK11 mutations we found distinct cellular patterns among LUAD and LUSC subtypes. Hence, our single-cell view of the TME provides further evidence for the link between the genetic make-up of the tumor, the histology, and the respective immune contexture (Wellenstein and de Visser, 2018).

**Figure 4.**
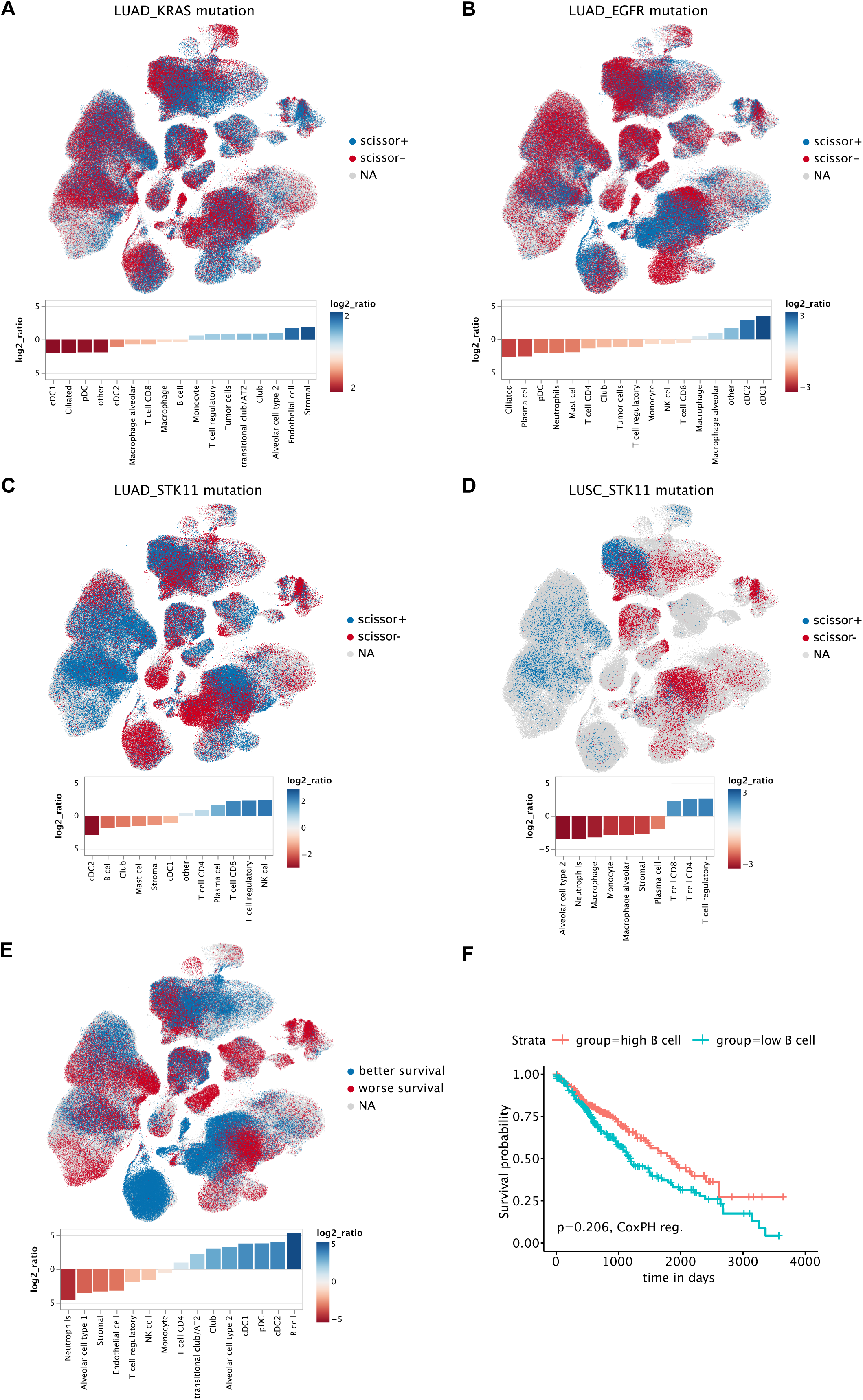
Association of cellular composition and distinct genotypes and survival in the TCGA data. (A-E) *SCISSOR* analysis relating phenotypic information from bulk RNA-seq data from TCGA with single cells. UMAP plots indicate the position of cells positively (blue) or negatively (red) associated with mutation or better survival. Bar charts indicate the mean log2 ratio of positively vs. negatively associated cells for each cell-type across all patients. A log2 ratio > 0 indicates a positive association with mutation, or better survival, respectively. Bars are only shown for cell-types with a log2 ratio significantly different from 0 at an FDR < 0.01 (paired Wilcoxon test). (A-C) Association of cellular composition with KRAS, EGFR and STK11 mutation in LUAD patients. (D) Association of cellular composition with STK11 mutation in LUSC patients. (E) Association of cellular composition with overall survival. (F) Kaplan-Meyer plot of patients with high (top 25%) and low (bottom 25%) B cell fractions of TCGA lung cancer patients as determined by deconvolution with EPIC. P-value has been determined using CoxPH-regression using tumor stage and age as covariates.

We next analyzed the cell type transcriptomic signatures and their association with survival in 1026 patients from the TCGA cohort. Overall, NSCLC patients with B cell-rich tumors showed a prominent association with improved survival, whereas neutrophils were the strongest negative predictor of survival (Figure 4E), and are the only immune cell type that was a negative predictor in both, LUAD and LUSC (Figure 4E, Figure S4C-D). Interestingly, other prognostically relevant cell types differed among LUAD and LUSC. For example, alveolar cell type II, plasma cells, cDC2s, and macrophages (including macrophages alveolar) associate with improved prognosis in LUAD, but are of poor prognosis in LUSC (Figure S4C-D). To support our finding, we used an independent method based on deconvolution using bulk RNA-seq data and confirmed that B cells are indeed associated with better prognosis, albeit significantly only for LUAD (Figure 4F, Figure S4E-F), which has also been proposed in multiple previous studies (reviewed in (Patel et al., 2020)). Similarly, the associations of neutrophils with poor outcome in many cancer entities including NSCLC has been previously shown with the deconvolution tool CIBERSORT (Gentles et al., 2015) and has also been reported by intratumoral measures of the neutrophil-to-lymphocyte ratio (Kargl et al., 2019) or blood measures (Alessi et al., 2021) of the neutrophil-to-lymphocyte ratio, respectively.

### TRNs acquire new properties in the TME

In a recent study remarkable neutrophil adaptability to different tissue environments was shown (Ballesteros et al., 2020), suggesting that while transient, TRN acquire new properties and function within tissue. To the best of our knowledge, comprehensive characterization of TRN in human NSCLC including both, TANs and NANs has not been carried out so far. TANs are known as very heterogenous cell population with both anti-tumorigenic and pro-tumorigenic properties. Previously suggested nomenclatures like N1 and N2 neutrophils most likely do not appropriately cover the diversity of neutrophil phenotypes and function (Jaillon et al., 2020). Likely due to technical reasons, the characterization of NANs lags even further behind that of TANs. To overcome the hitherto insufficient TRN characterization, we here characterized the transcriptomic signatures of TANs and NANs in NSCLC using the extended atlas (Figure 5A-B). Neutrophils were more abundant in LUSC compared to LUAD patients (Figure S5A). Flow cytometry analysis of 47 LUAD and 16 LUSC patients confirmed a significant increase in neutrophils infiltration in LUSC tumors (Figure 5C). The overall TAN phenotype was characterized by high expression of *OLR1* (LOX-1)*, VEGFA, CD83,* and *CXCR4,* and low expression of *CXCR1, CXCR2, PTGS2*, *SELL* (CD62L)*, CSF3R* and *FCGR3B* (CD16B) (Figure 5D). The TAN characteristic gene set included expression patterns of both, established neutrophil markers (*CXCR1, CXCR2, CXCR4, PTGS2*) as well as novel candidates (*OLR1, VEGFA, CD83*) as discussed below.

**Figure 5.**
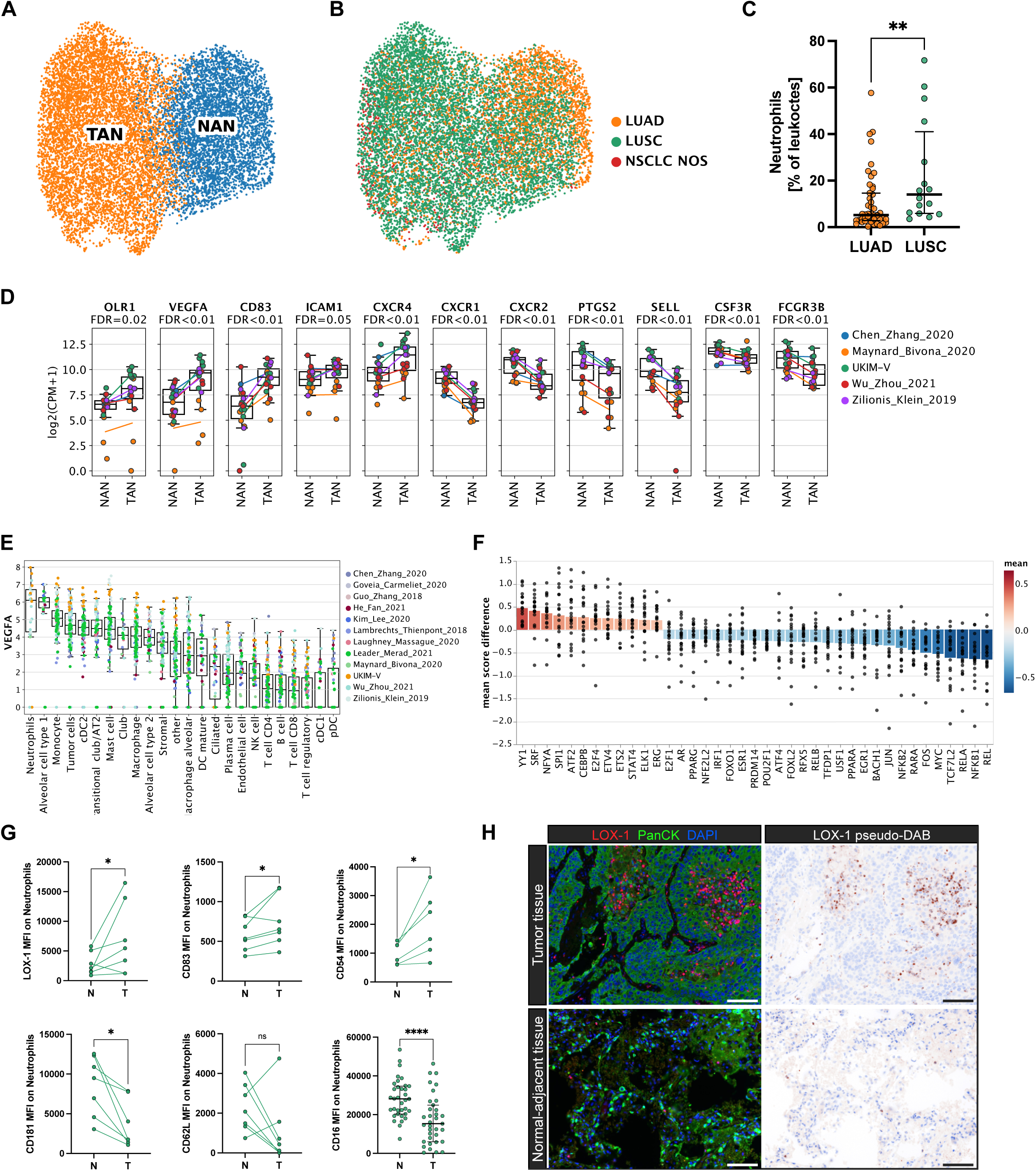
Characterization of tissue-associated neutrophils using scRNA-seq. (A-B) UMAP of tissue-resident neutrophils (TRN) from the extended atlas, (A) classified into tumor-associated neutrophils (TANs) and normal-adjacent associated neutrophils (NANs), and (B) colored by histology. (C) Neutrophil fractions (as percentage of leucocytes) by flow cytometry of LUAD and LUSC tumor tissue (LUAD n=47, LUSC n=16; Wilcoxon test, **p<0.01). The horizontal line represents the median, whiskers extend to the inter-quartile range. (D) Candidate TAN genes. Each dot refers to a patient with at least 10 neutrophils in both NAN and TAN groups. Lines indicate the mean per study. P-values are derived from a paired t-test and adjusted for false-discovery rate (FDR). (E) Analysis of the expression levels of VEGFA in various cell-types in primary tumor samples. Each dot represents a patient with at least 10 cells, colored by study. (D-E) In all boxplots, the central line denotes the median. Boxes represent the interquartile range (IQR) of the data, whiskers extend to the most extreme data points within 1.5 times the IQR. (F) Transcription factor analysis of TANs using DoRothEA. The y axis indicates the difference from the mean transcription factor activity score between groups. Each dot represents a single patient, bars the mean of all patients. P-values are derived using a paired t-test and FDR-adjusted. The bar chart only shows transcription factors with a mean score difference > 0.2 and a FDR < 0.1. (G) Comparison between tumor and normal-adjacent samples for selected candidate genes using flow cytometry. Each dot represents a patient that was not part of the scRNA-seq dataset. Paired Wilcoxon test, *p<0.05, ****p<0.0001. CD16: the horizontal line represents the median, whiskers extend to the inter-quartile range. (H) Selected multiplex immunofluorescence (M-IF) stainging of LOX-1 (red) and pancytokeratin (green) in tumor tissue and matched normal adjacent lung tissue of a patient with LUSC. Scale bar = 100 µm.

The neutrophil phenotype differs in dependence of spatial-, temporal- and disease-specific clues (Hedrick and Malanchi, 2022) as well as during the evolution from bone marrow-resident immature (CXCR4^high^, CXCR2^low^, CD16^low^, CD62L^low^, MME^low^), to circulating-mature (CXCR2^high^, CD16^high^, CD62^high^), to aged/senescent neutrophils (CXCR4^high^) (Coffelt et al., 2016; Evrard et al., 2018; Martin et al., 2003). However, none of these markers is specific for a certain maturation state. Relative to TANs, matched NANs in our dataset showed high expression of established neutrophil maturity markers (*SELL, PTSG2, CXCR2, CXCR1, FCGR3B, MME*) as well as canonical neutrophil markers (*S100A8, S100A9*) (Figure S5B). While downregulation of these markers in TANs suggests immaturity, we could not identify a clear expression pattern of previously suggested immaturity signatures (Evrard et al., 2018; Schulte-Schrepping et al., 2020). Notably, *CXCR1* and *CXCR2* have previously been reported as TAN markers in NSCLC (Wu et al., 2021; Zilionis et al., 2019), however our analysis revealed elevated *CXCR1*/*CXCR2* expression in NANs. One explanation for this discrepancy could be that these studies lack matched tumor adjacent tissue to correlate their findings. Conversely, low expression of *SELL* (CD62L) and *CXCR2* (both downregulated in aged neutrophils (Lin et al., 2020)) and high expression of the known neutrophil activation markers *CD83* (an inhibitory immune checkpoint) (Li et al., 2019; Yamashiro et al., 2000), the atypical chemokine receptor *CCRL2* (Del Prete et al., 2017), *ICAM1* (CD54) (Lin et al., 2020)*, C15orf48* (a mitochondrial transcript upregulated during inflammation) (Clayton et al., 2021) as well as several cytokines (*CCL3, CCL4, CXCL2)* (Figure S5B) support an aged/chronically activated/exhausted TAN phenotype (Zhang et al., 2015).

Neutrophils support the pro-angiogenic switch in cancer *via* release of VEGF and other pro-angiogenic factors (reviewed in (Ozel et al., 2022)). Remarkably, our atlas provided evidence that neutrophils represent a major source for *VEGFA* expression within the NSCLC TME (Figure 5E). A highly differentially expressed TAN marker of major interest is lectin-type oxidized LDL receptor 1 (LOX-1) encoded by the *OLR1* gene that is known as main receptor for oxidized low-density lipoprotein (LDL) (Gonzalez-Chavarria et al., 2014) (Figure 5D). *OLR1* has been described as putative marker to distinguish normal peripheral blood neutrophils (LOX-1^-^) from polymorphonuclear myeloid-derived suppressor cells (PMN-MDSC) (Condamine et al., 2016), respectively. Currently, a clear distinction of PMN-MDSCs and TANs is not feasible due to the lack of cell-type specific markers and direct comparative analysis, thus, it remains unclear whether these are distinct cell types. However, concordant to our results, *OLR1* expression by TANs has been previously described (Wu et al., 2021) and our comparative analysis to matched NANs underlined the tumor-specificity of this marker. Moreover, peroxisome proliferator-activated receptor gamma (PPARG), a nuclear receptor and direct transcriptional regulator of *OLR1* (Chui et al., 2005), was elevated in TANs (Figure 5F). Concordantly, flow cytometry analysis of tissue from NSCLC patients (n=7) confirmed elevated LOX-1 (OLR1) expression in TANs (Figure 5G). We could further validate the transcriptomic TAN signature at protein level by flow cytometry, including elevated expression of CD83 (n=7), CD54 (ICAM1) (n=6) and lower expression of CD181 (CXCR1), CD62L (SELL) (n=8), and CD16 (n=33) (Figure 5G).

We additionally performed multiplex immunofluorescence staining of paraffin-embedded NSCLC tumor tissue and patient-matched normal adjacent lung tissue. Co-staining of LOX-1 and CXCR2 suggested LOX-1 as neutrophil marker (Figure S5C) (of note, CXCR2 also marks MDSCs (Steele et al., 2016)). We found infiltration of LOX-1 positive cells in tumor tissue but not in adjacent normal-lung tissue (Figure 5H), underlining the cancer tissue-specificity of this marker.

In summary, comprehensive characterization of TANs at single-cell level identified subsets with distinct phenotypes, and indicates the potential of TANs to acquire antigen presenting-like properties to elicit anti-tumor immunity.

### Plasticity and non-canonical functional properties of TRNs

Previous studies have proposed transcriptomic subclusters of neutrophils in NSCLC (Zilionis et al., 2019) and COVID-19 infection (Schulte-Schrepping et al., 2020). However, a distinct sub-classification and in-depth characterization of TRN including TANs and NANs has not been described so far. We applied unsupervised Leiden-clustering on all atlas neutrophils (n=12,467), separating four TAN subsets (TAN-1 to TAN-4) and two NAN subsets (NAN-1 and NAN-2) (Figure 6A) that are backed by multiple datasets (Figure S6A) and multiple patients (Figure S6B). Marker gene selection revealed an extensive phenotypic heterogeneity among the clusters and allowed identification of marker genes for each sub-cluster, of which the top 10 are given in Figure 6B. The TAN signature genes described above (*OLR1, VEGFA, CXCR4, CD83*) showed relative homogenous expression among all TAN subclusters (Figure S6D). Overall, NAN clusters showed a predominance in LUAD and TAN clusters in LUSC tumors (Figure 5B).

**Figure 6.**
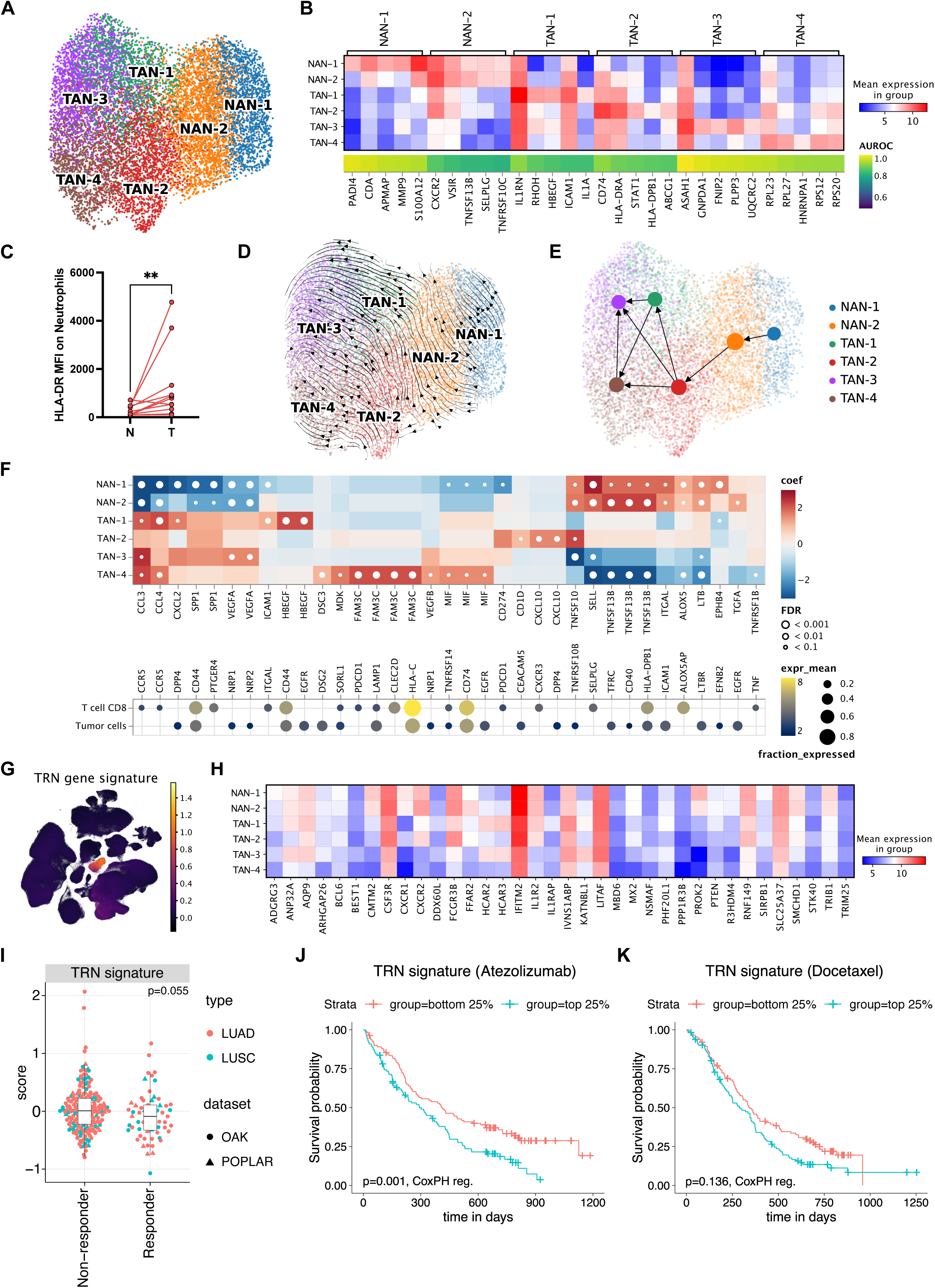
TAN subtypes in NSCLC. (A) UMAP of all neutrophils colored by TAN and NAN subclusters. The neutrophil clusters derive from 32 patients with each > 10 neutrophils. (B) Top 5 markers for each TAN and NAN cluster. The marker gene quality is reflected by the area under the receiver operator characteristics curve (AUROC, 1= marker gene perfectly distinguished the respective cluster from other clusters in all patients; AUROC 0.5 = no better than random) (C) Flow cytometry analysis of tumor and normal-adjacent tissue for HLA-DR. Each dot represents a patient. Paired Wilcoxon test, **p<0.01. (D) UMAP of neutrophils from the UKIM-V dataset with RNA velocitity vectors projected on top. (E) Partition-based graph abstraction (PAGA) based on RNA velocities, projected on the UMAP plot. Edges represent cell-type transitions called by PAGA. (F) Outgoing interactions of TRN subclusters with tumor cells and CD8^+^ T cells. Upper panel: Differentially expressed ligands in each subcluster (linear model on log-normalized pseudobulk samples with sum-to-zero-coding, FDR < 0.1, abs. log2 fold change > 1). Heatmap colors indicate log2 fold changes clipped at ±3, where red and blue indicate up- and downregulation the the corresponding cluster, respectively. Lower panel: Respective receptors and the expression by cell type. Dot sizes and colors refers to the fraction of cells expressing the receptor and gene expression, respectively, averaged over all patients. Dots are only shown for receptors that are expressed in at least 10% of the respective cell-types. (G) UMAP of the extended atlas colored by the score of the TRN gene signature (36 genes with high specificity for TRNs) (H) Heat map of the TRN gene signature across the TRN subclusters. Colors indicate the mean gene expression across patients in the respective clusters. (I-K) Predictive value of the TRN signature in bulk RNA-seq data from the OAK (Rittmeyer et al., 2017) and POPLAR (Fehrenbacher et al., 2016) cohorts of NSCLC patients treated with atezolizumab (anti-PD-L1) or docetaxel (chemotherapy). (I) Comparison of non-responders (progressive disease) with responders (complete response, partial response) treated with atezolizumab, shown for each histotype. The p-value has been determined using logistic regression, including the dataset and tumor type as covariates. (J) Kaplan-Meyer plot comparing patients treated with atezolizumab with high (top 25%) and low (bottom 25%) TRN signature scores. P-value has been determined using CoxPH-regression including cohort and histology as covariates. (J) Kaplan-Meyer plot comparing patients treated docetaxel with high (top 25%) and low (bottom 25%) TRN signature scores. P-value has been determined using CoxPH-regression including cohort and histology as covariates.

Specific NAN-1 genes included the alarmin *S100A12*, a known marker of activated proinflammatory neutrophils (Meijer et al., 2012), the NETosis co-factor *PADI4* (Leshner et al., 2012) as well as pro-angiogenic markers (*PROK2*, *MMP9*). The NAN-2 cluster is similar to the NAN-1 cluster, but showed reduced expression of some NAN-1 genes (e.g. *S100A12*, *MME*, *PROK2*) (Figure S6C). TAN-1 was characterized by high expression of *CXCL8*, a core CXCR2 and CXCR1 agonist that mediates neutrophil activation and neutrophil extracellular trap (NET) formation (Ha et al., 2017), as well as by genes that regulate neutrophil activation, recruitment or cell adhesion such as *CXCL1*, *CXCL2* (Girbl et al., 2018; Lentini et al., 2020), *ICAM1* (Yang et al., 2005), and *CD44* (Katayama et al., 2005). These finding support the concept that the plasticity of neutrophils is shaped by the NSCLC TME that attracts and activates neutrophils.

The TAN-2 subcluster was characterized by the expression of the MHC II genes *HLA-DRA, CD74* and *HLA-DPB1*, indicating a phenotype with immunogenic antigen presenting feature. Both *CD74* and *HLA-DRA* are also expressed in the other TAN clusters albeit at lower level (Figure 6B). This observation implies acquisition of antigen presenting-like properties by neutrophils at the tumor site, as previously reported (Singhal et al., 2016). Such conversion of neutrophils to antigen presenting cells may elicit anti-tumor immunity and has recently been shown in a murine model (Mysore et al., 2021). To validate the scRNA-seq findings, we analyzed samples from additional 11 patients using flow cytometry, confirming the antigen presenting phenotype as seen by an up-regulation of HLA-DR on TANs in NSCLC compared to normal adjacent lung tissue (Figure 6C). Of note, transition to HLA-DR^+^ neutrophils was accompanied by a shift towards the identified TAN signature (elevated expression of CD83, LOX-1 and lower expression of CD181 (CXCR1), CD62L (SELL) and CD16) in neutrophils derived from NSCLC tumor tissue (Figure S6F). Also, TAN-2 and NAN-2 clusters showed high expression of interferon-associated genes (*IFIT1, IFIT2, IFIT3, XAF1, ISG15*) (Figure S6C), a signature that has previously been described for non-activated mature neutrophils in COVID-19 (Schulte-Schrepping et al., 2020) as well as in NSCLC (Zilionis et al., 2019).

The TAN-3 cluster was characterized by a high expression of the lipid metabolism-related genes *PLIN2* (allowing degradation of intracellular lipid droplets as important metabolic pathway) and *PLPP3* (reported to act as negative regulator of inflammatory cytokines) (Touat-Hamici et al., 2016). Of note, neutrophils may transfer lipids to cancer cells, enhancing their proliferation (reviewed in (Hedrick and Malanchi, 2022)). TAN-3 was characterized by a high expression of *MAP1LC3B* (LC3) that marks autophagy, a critical lipolytic process providing energy for neutrophil differentiation and function (Riffelmacher et al., 2017). Also, TAN-3 showed high expression of *PLAU*, encoding the plasminogen-activator urokinase (uPA) that is known to activate extracellular matrix–degrading proteases and to induce intracellular signaling pathways by binding its cognate receptor uPAR, thus regulating cell adhesion and (tumor cell) migration. It has previously been reported that uPA and uPAR are stored in primary neutrophil granula (Pedersen et al., 2000), while uPAR is expressed by cancer cells (i.e. in response to hypoxia) (Smith and Marshall, 2010). Thus, TAN-3 may be involved in cancer cell metastasis formation. Finally, TAN-4 showed high expression of ribosomal genes (such as *RPL10, RPS2, RPS18, RPL3*), similar to a neutrophil cluster identified in severe COVID-19 patients (Schulte-Schrepping et al., 2020), and could suggest a plastic phenotype of these TAN-4 transitioning to another cell phenotype, as described previously for tumor endothelial cells (Goveia et al., 2020).

The transcriptional profiles of the neutrophil subsets indicate their remarkable phenotypic plasticity. We therefore performed RNA velocity analysis (see Methods) using only the UKIM-V datasets since the method requires raw sequencing data. The analysis indicates transition from NAN-1 to NAN-2 and then from NAN-2 to TAN-2, that further segregates into TAN-1, TAN-3, and TAN-4 (see Figure 6D and partition-based graph abstraction (Wolf et al., 2019) in Figure 6E). Interestingly, this TRN evolution seems to follow a one-directory path with TAN-3 as final transition, although at this point we do not know whether TAN-3 could further transit to a different cell type. These observation supports the hypothesis that TAN phenotypes are modulated by local cues encountered in the TME (Shaul and Fridlender, 2019).

We next investigated the cellular interactions of TRN subsets with CD8^+^ T cells and tumor cells by analyzing differentially expressed TRN ligands (FDR<0.1, abs(log2FC)>1) of each subset, revealing distinct signaling of NANs versus TANs (Figure 6F). In all TAN subsets we found high expression of VEGFA, again underlining their important proangiogenic role, as well as SPP1 signaling that has previously been associated with immunosuppressive TME compartments (Zheng et al., 2021) and pro-EMT effects (Zhao et al., 2018). We also found TRN subset specific signaling patterns, including high *FAM3C* and *MIF* expression by TAN-4, genes that have been attributed to pro-metastatic functions (Funamizu et al., 2013; Noguchi et al., 2018), as well as high CXCL10-CXCR3 signaling of TAN-2 to CD8^+^ T cell, suggesting chemotactic effects of this subset. Conversely, NANs showed prominent interactions involving genes of the tumor-necrosis family (*TNFSF13B*, *TNFSF10*, *LTB*) that have been previously associated with neutrophil activation (Afonso et al., 2012; Besteman et al., 2020).

Overall, the analyses of the neutrophil compartment provides evidence for the high diversity and plasticity of TRNs in NSCLC, suggesting a major impact on clinical outcomes of patients treated with checkpoint-inhibitors.

### TRN gene signature is associated with immune checkpoint inhibitor treatment failure

Our deconvolution of the diversity of TRNs at the single-cell level prompted us to relate this information to patient prognosis and response to both, chemotherapy and therapy with immune checkpoint inhibitors. Using common cut-off for finding specific marker genes for cell-type estimation from bulk RNA-seq samples ((Becht et al., 2016), see methods section), we derived a signature of 36 genes that are highly specific for TRNs compared to all other cell types based on the single-cell data from the primary tumor and the normal adjacent tissue (Figure 6G). By intersecting the TRN signature with the TAN and NAN markers from above, we identified only a small number of genes that are specific for TANs (*KATNBL1*, *PI3*), and NANs (*CMTM2*, *PROK2*, *CXCR1*, *CXCR2*, and *FCGR3B*) and expressed only at a very level in other cells. The expression of the TRN signature genes was heterogeneous between the different TRN subsets (Figure 6H). In order to analyze the prognostic and predictive value of the TRN signature, we used bulk RNA-seq data from pre-treatment tumors from POPLAR (Fehrenbacher et al., 2016) and OAK (Rittmeyer et al., 2017), two randomized clinical trials of anti-PD-L1 antibody (atezolizumab) versus chemotherapy (docetaxel) in NSCLC, representing the largest transcriptional collection in these settings (Patil et al., 2022). In total there were 891 patients of which 439 were treated with atezolizumab (316 LUAD and 123 LUSC) and 452 with docetaxel (312 LUAD and 139 LUSC). The TRN gene signature was associated with anti-PD-L1 therapy failure in these NSCLC cohorts (Figure 6I). Analyses of the survival data from these cohorts showed that the prognostic benefit of the TRN signature was significant for the immunotherapy arm (Figure 6J) but not for the chemotherapy arm (Figure 6K). The prognostic value for the immunotherapy arm was stronger for LUSC (Figure S6G) compared to LUAD (Figure S6H). We then analyzed the lung cancer immune activation module (LCAM) which confers prognostic benefit in anti-PD-L1 treatment but not for chemotherapy (Leader et al., 2021). The low correlation between the TRN signature and the LCAM (Figure S6I) suggests that the TRN are not associated with frequencies of the lineages defined by the LCAM (T_activated_, IgG^+^ plasma cells, and SPP1^+^ macrophages (Leader et al., 2021)), and indicates that the TRN signature is an independent marker.

## DISCUSSION

We built a large-scale atlas of single-cell transcriptomes of NSCLC through integration of 21 studies spanning over 1,200,000 cells from 538 samples and 309 individuals representing 1.62 billion expression values. Reduction of dataset-specific batch effects due to variation in experimental design and used platforms while retaining biological information resulted in a high-quality reference atlas for NSCLC, offering superior coverage of histological and clinical variables and thereby providing a unique resource for dissecting the cellular diversity in the TME. We leveraged the information content of the NSCLC atlas by sequencing additional patient samples using a platform suitable to depict low-mRNA content cells, which enabled us to comprehensively characterize the diversity and plasticity of TRNs. The analysis of the NSCLC atlas and the integration of large-scale bulk RNA-seq dataset provided several important novel biological insights.

First, we provide a high-resolution view of the TME in NSCLC with 38 major cell types/states and show different cell type composition patterns in LUAD and LUSC including more precise classification of epithelial cells in both histotypes. The single-cell composition of the TME in NSCLC enabled refined tumor classification and patient stratification into four immune phenotypes: immune deserted, myeloid, B cell, and T cell subtypes. The myeloid, B cell, and T cell subtypes showed predominantly LUAD histology whereas half of the LUSC patients were assigned to the immune deserted subtype. Moreover, the four tumor immune phenotypes showed different pathway activation and cytokine expression in tumor cells, implicating that the hetero-cellular crosstalk is determined by histotypes and tumor cell states. We also provide evidence that the hetero-cellular crosstalk patterns are associated with distinct tumor immune phenotypes.

Second, integration of bulk RNA-seq data from the TCGA NSCLC cohort with our single-cell atlas enabled identification of cell subpopulations driving the phenotype of interest. We uncovered cell subsets associated with alterations in major driver genes, such as *EGFR*, *KRAS*, *STK11*, and *TP53* in both, LUAD and LUSC subtypes, providing further evidence that genetic aberrations in cancer cells dictate the immune contexture of tumors (Wellenstein and de Visser, 2018). Previous genotype-immunophenotype studies in human cancers focused on mutational load or particular driver genes and major immune cell types (Wellenstein and de Visser, 2018). Here, we provide a high-resolution genotype-immunophenotype mapping, and thereby generated hypotheses that could lead to mechanistic insights into the causal relationships between tumor genetics and immune cell composition. Moreover, this integration of large-scale single-cell atlas with large-scale bulk data sets (TCGA) enables the identification of cell subpopulations that drive other phenotypes such as disease stage, treatment response, and survival outcome. For example, we show that B cells were associated with better survival whereas neutrophils are associated with poor outcome in NSCLC patients. Integration of other bulk datasets is straightforward and could help to identify biologically and clinically relevant cell subpopulations for the phenotype of interest.

Third, since neutrophils are underrepresented in most scRNA-seq studies, we generated data using scRNA-seq platform capturing also low-mRNA content cells and provide for the first time in-depth characterization of TRNs including both, TANs and NANs in human NSCLC. We show that TRN acquire new properties and function in the TME and that the TAN phenotypes are characterized by both established neutrophil markers (*CXCR1, CXCR2, CXCR4, PTGS2*) as well as novel candidates (*OLR1, VEGFA, CD83*). Over and above, in-depth analyses of the TRN transcriptomes revealed extensive phenotypic heterogeneity among the TAN and NAN subsets. TANs are an important component of the TME and can exert dual functions (Jaillon et al., 2020). They may either alter the TME, e.g. by influencing adaptative immune responses, promoting angiogenesis (Bekes et al., 2011; Deryugina et al., 2014; Nozawa et al., 2006), and by extracellular matrix remodeling with formation of extracellular traps to shield cancer cells from cytotoxicity (Teijeira et al., 2020). Conversely, TANs can mediate antitumor responses by directly inducing cancer cell DNA damage/genomic instability via reactive oxygen species (ROS) release (Knaapen et al., 1999; Wculek et al., 2020). Conflicting evidence indicates both pro-and anti-tumor properties, such as the dichotomous interaction of TANs with effector T lymphocytes – TANs may be pro-inflammatory by activating T cells through antigen presentation, but have also been described as immunosuppressive by secretion of arginase-1 and ROS. This latter immunosuppressive phenotype has also been postulated as cancer-specific low-density granulocytic myeloid derived suppressor cells (G-MDSC) (Jaillon et al., 2020). Our dissection of the diversity of TANs suggest that the conflicting reports can be attributed to the different TRN subsets. Of particular interest for cancer immunotherapy is the TAN phenotype with immunogenic antigen-presenting feature. Identification of targets that can block the transition of the antigen-presenting TAN-2 subset into TAN-3 and TAN-4 or reverse the final phenotypes into TAN-2 phenotype is an important goal for future studies.

Finally, we report that TRN-derived gene signature has a predictive and prognostic effect of the TRN signals for immunotherapy treated NSCLC patients. It has been shown previously that the ratio of CD8^+^ T cells to neutrophils in the TME of NSCLC was capable of separating the patients responsive to immunotherapy from those with stable or progressive disease (Kargl et al., 2019). However, only a small cohort (n=28) treated with both anti-PD1 (pembrolizumab, nivolumab) and PD-L1 antibodies (atezolizumab, durvulamab) was used. Here, using the largest available transcriptomic data for NSCLC patients (n=439) from two randomized clinical trial cohorts treated with a single anti-PD-L1 antibody (atezolizumab), we provide an evidence for the association of TRNs with therapy failure. Notably, lack of association with previously reported LCAM module (Leader et al., 2021), indicates that the effect is independent of the cell types that define the LCMA module (T_activated_, IgG^+^ plasma cells, and SPP1^+^ macrophages).

Beyond these biological insights, the results from this study have also important implications. Specifically, the diversity and plasticity of TRNs shown here further underscores the necessity to reevaluate the rationale for targeting neutrophils to overcome immune checkpoint inhibitor therapy resistance in combination therapies using CXCR1 and CXCR2 antagonists and other inhibitors (Zhang and Houghton, 2021). As shown here, TANs can acquire antigen presenting properties and such conversion of abundant neutrophils to antigen-presenting cells could overcome the limitations of the low abundance of cross-presenting DCs (Mysore et al., 2021). We advocate that rigorous approaches are required to analyze the impact of the TRN diversity and plasticity on tumor immunity in NSCLC and possibly in other cancers. Moreover, this finding is not restricted to cancer studies. For example, in SARS-CoV2 infection, neutrophils act as a major inflammatory driver and they constitute up to 70% of leukocytes in bronchoalveolar lavage fluid of patients with COVID-19 pneumonia (Dentone et al., 2021). Yet, many of the published scRNA-seq analysis of COVID-19 patient samples (e.g. lung tissue, peripheral blood, bronchoalveolar lavage) did not dissect a significant neutrophil population among the myeloid cell compartment (Delorey et al., 2021; Ren et al., 2021). Hence, it is likely that the underrepresentation or even absence of neutrophils in most transcriptomic analysis does not reflect the actual cellular composition and can be ascribed to technical issues that hinder appropriate capturing.

In conclusion, our NSCLC atlas provides a single-cell resource and represents an important contribution to the community. We also provide web a portal that enables interactive exploration of the dataset through cell-x-gene (https://luca.icbi.at), a web-based viewer for single-cell datasets that allows visualization of metadata and gene expression. Users can project their own datasets onto the core atlas using scArches (Lotfollahi et al., 2022) and perform automated cell-type annotation, thereby enabling detailed annotation of new datasets and identification of rare cell identities. Thus, the biological insights we present here and future discoveries arising from the exploitation of the high-resolution NSCLC atlas could provide the basis for developing combination therapies for NSCLC patients who are not sufficiently responding to immune checkpoint blockers.

## ACKNOWLEDGMENTS

This work was supported by the European Research Council (grant agreement No 786295 to ZT), and by the Austrian Science Fund (FWF) (project I3978 to ZT). GS was supported by a DOC-fellowship from the Austrian Academy of Sciences. ZT is a member of the German Research Foundation (DFG) project TRR 241(INF). DW work was supported by the “Deutsche Krebshilfe” (grant No. DKH 70112994). FF was supported by the Austrian Science Fund (FWF) (T 974-B30) and by the Oesterreichische Nationalbank (OeNB) (18496). DW, SiS and AP were supported by the In Memoriam Gabriel Salzner Stiftung. SiS was supported by the FFG grant Austrian Research Promotion Agency (858057 HD FACS) and LH by the OEGHO Förderpreis Onkologie 2021. The results published here are in part based upon data generated by the TCGA Research Network: https://www.cancer.gov/tcga. The authors would like to thank Marcus Kalb, Sophia Daum, Annabella Pittl and Elisabeth Hoflehner for technical support.

## AUTHOR CONTRIBUTIONS

Conceptualization, GS, StS, LH, DW, AP, and ZT; data analysis, GS, GF, EP, FF, LH, StS, SiS and DR; human samples, LH, GG, KS, FA, GP, StS, AP, DW; single-cell sequencing, StS, GU, MS; sample analysis, StS, SiS, MS, GU; writing-original draft, LH, StS, and ZT; writing - review and editing, all authors; funding acquisition: DW, SiS, AP, and ZT.

## SUPPLEMENTARY INFORMATION

Figures S1-S7

Tables S1-7

## Key Resource Table

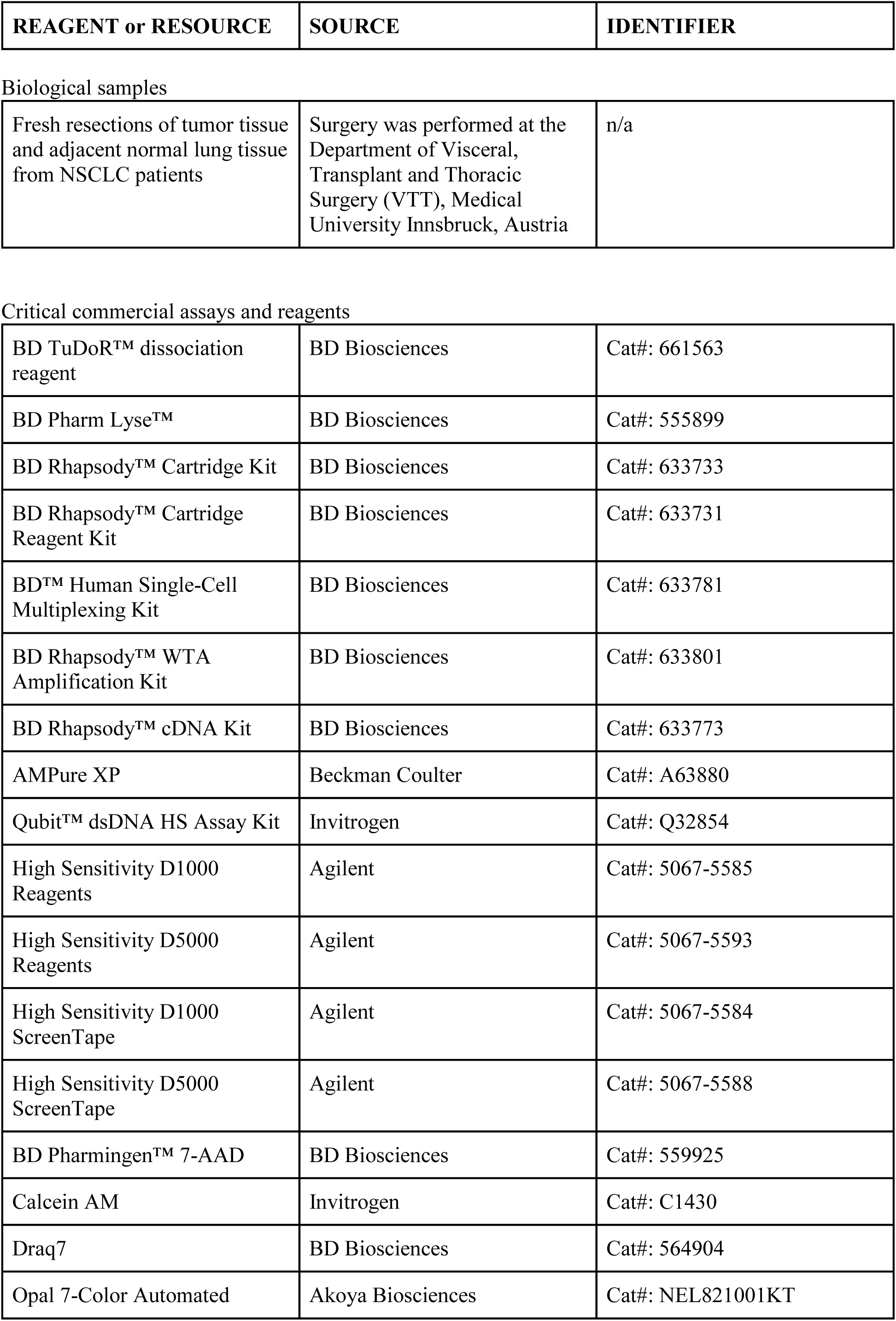
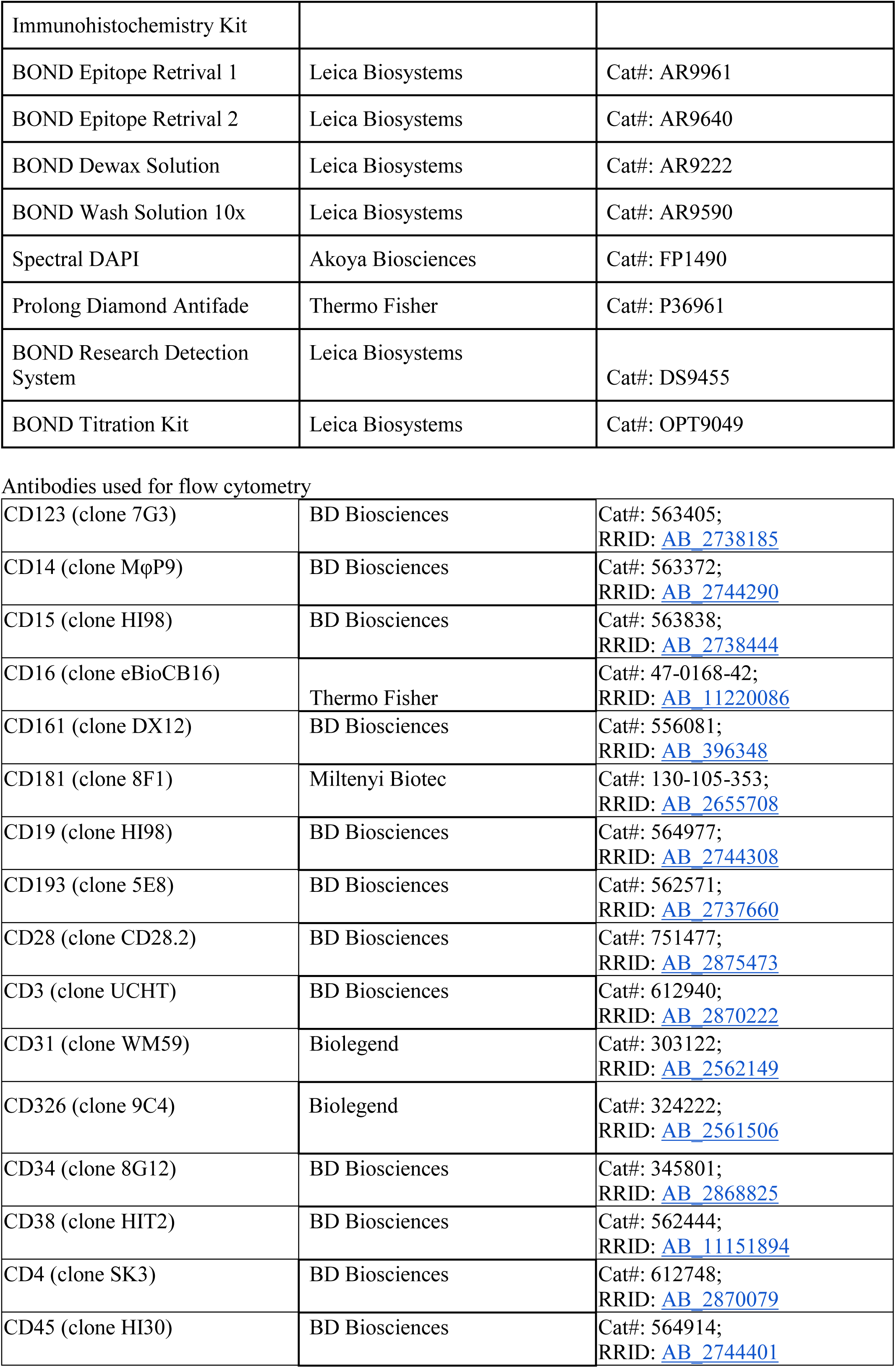
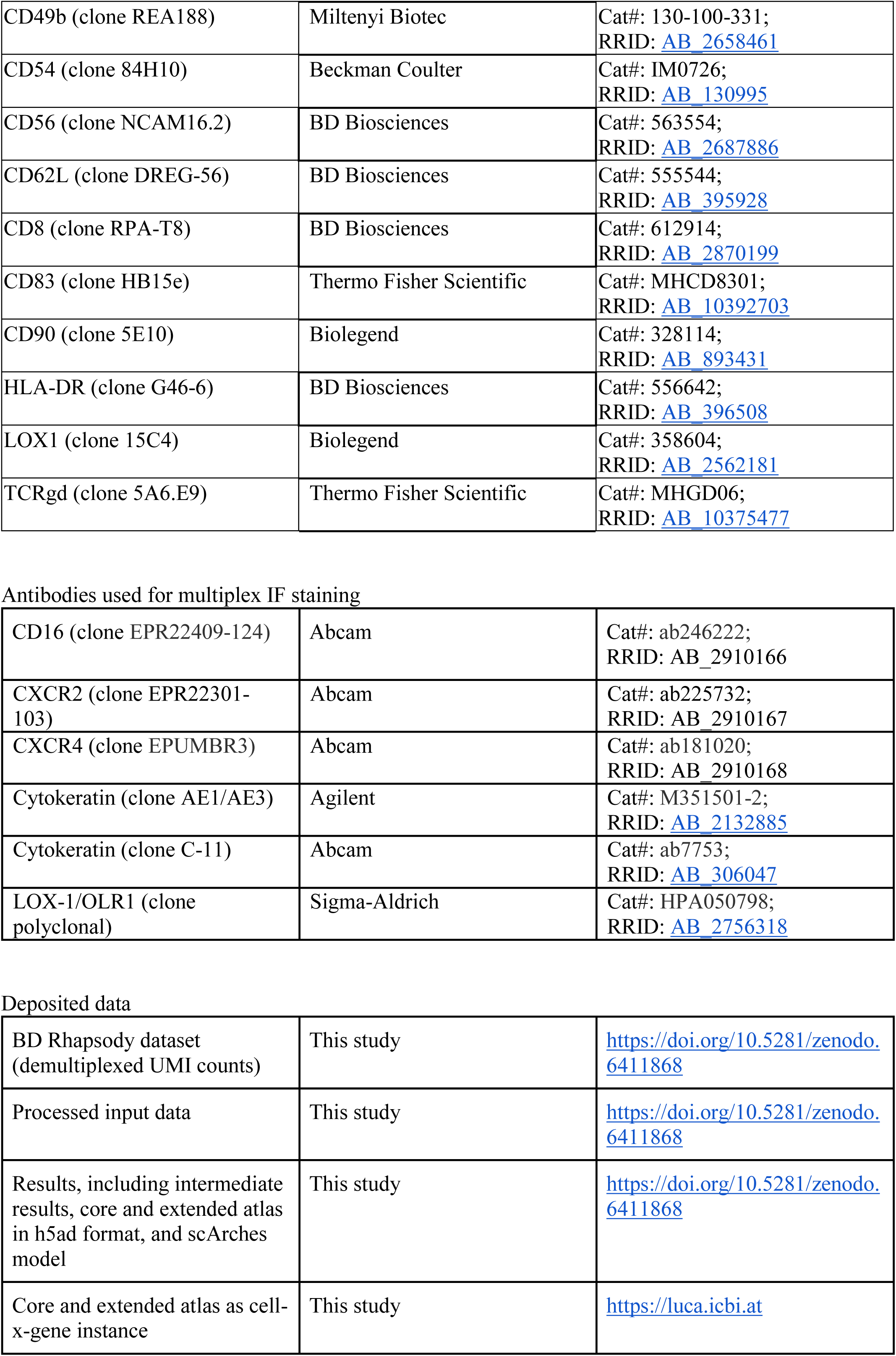
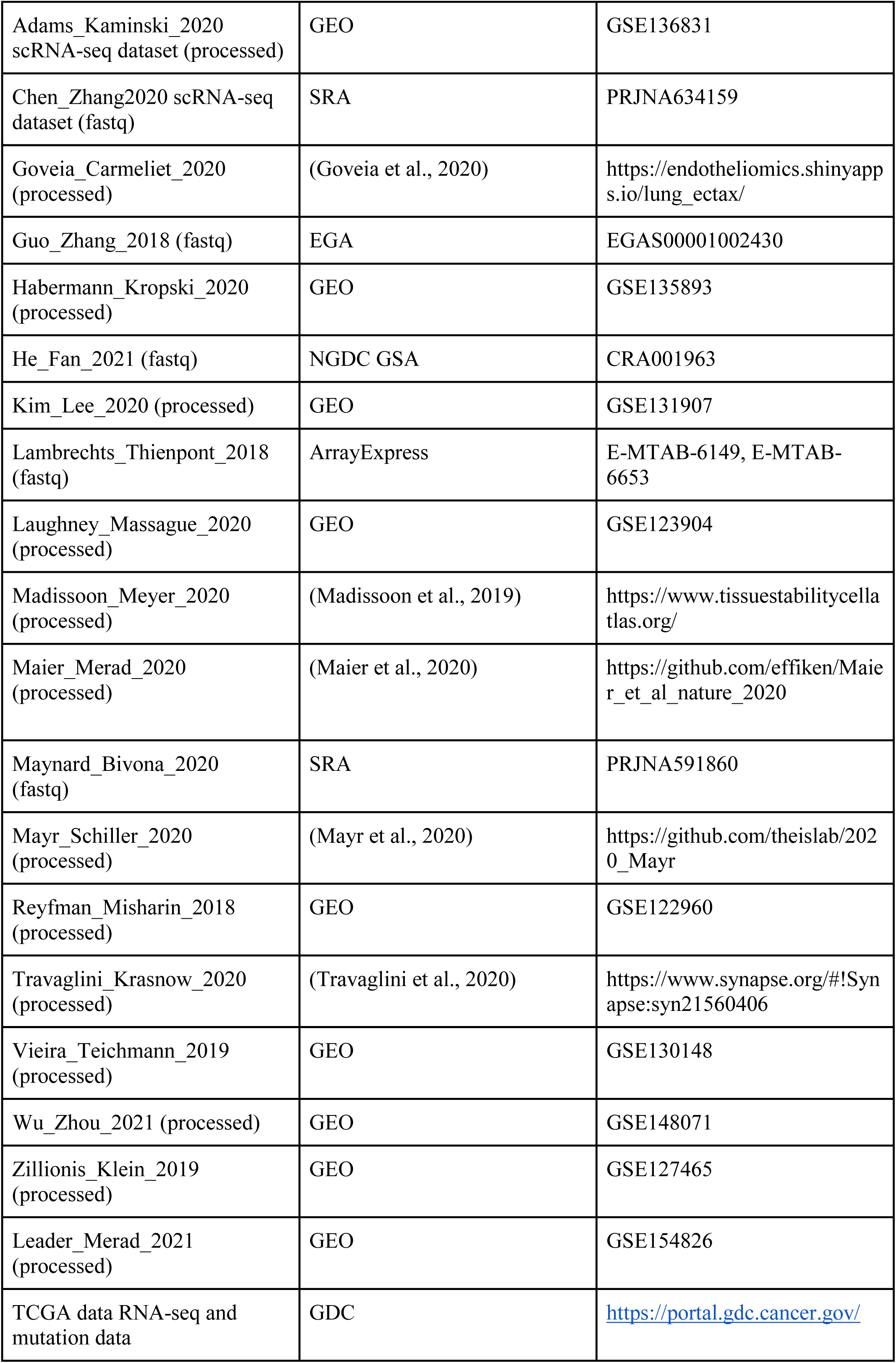
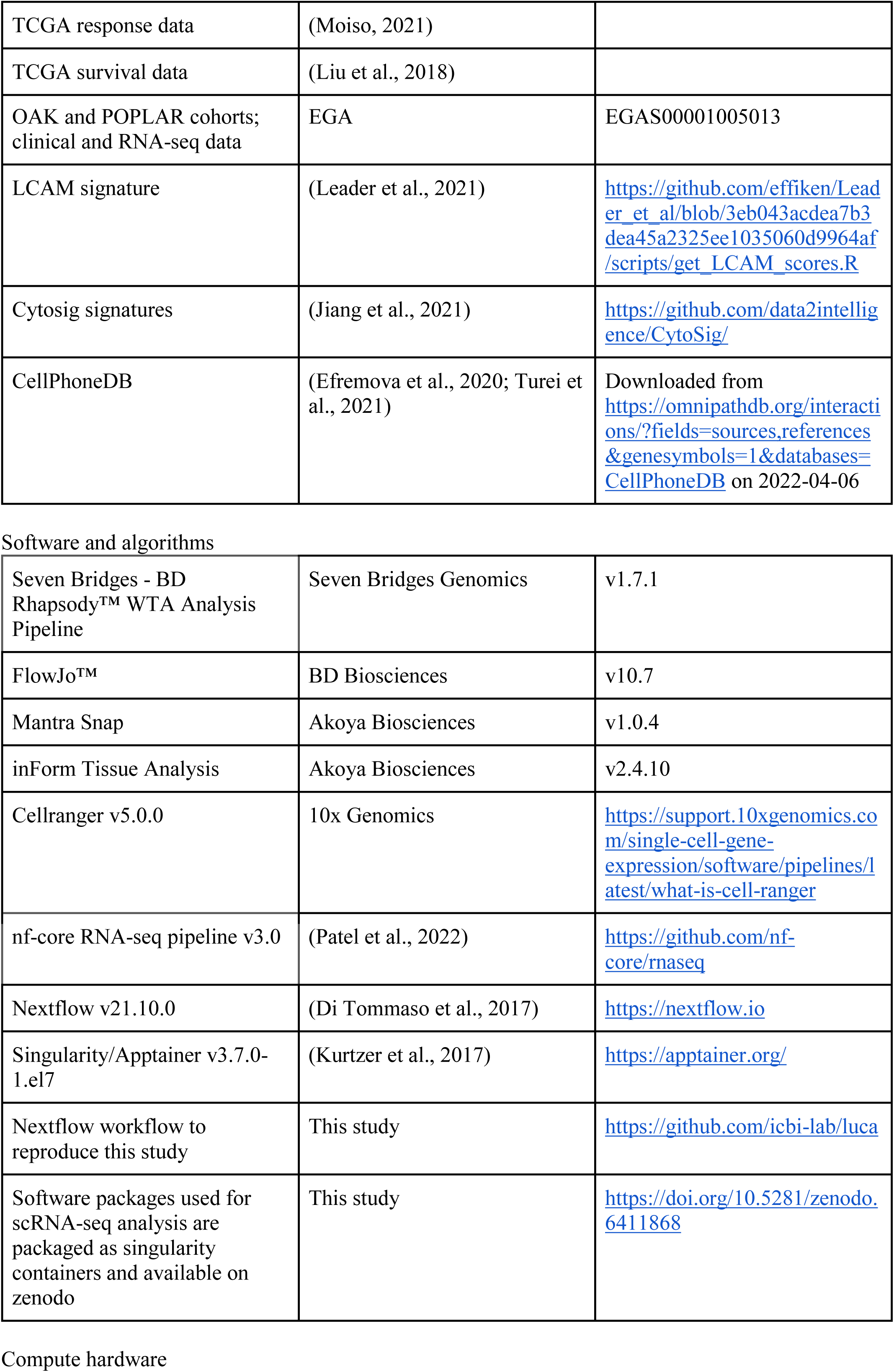
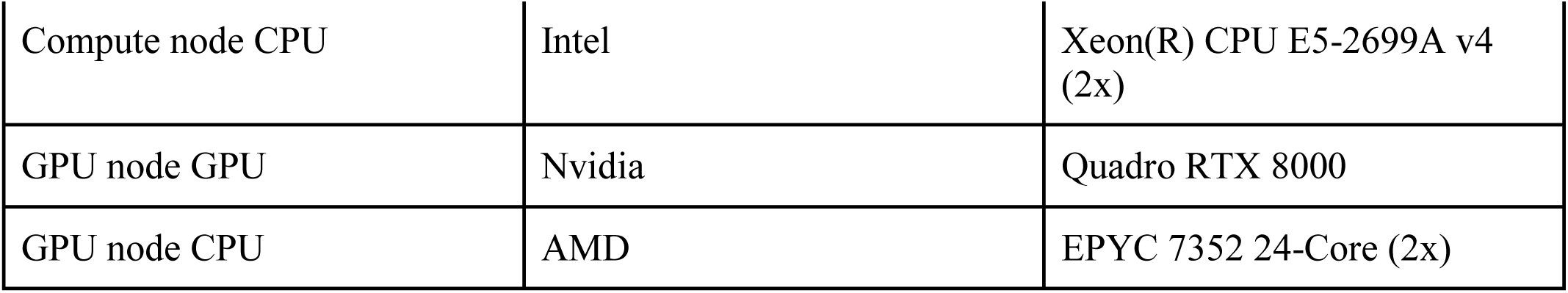

| REAGENT or RESOURCE | SOURCE | IDENTIFIER |
| --- | --- | --- |
| Biological samples |  |  |
| Fresh resections of tumor tissue and adjacent normal lung tissue from NSCLC patients | Surgery was performed at the Department of Visceral, Transplant and Thoracic Surgery (VTT), Medical University Innsbruck, Austria | n/a |
| Critical commercial assays and reagents |  |  |
| BD TuDoR™ dissociation reagent | BD Biosciences | Cat#: 661563 |
| BD Pharm Lyse™ | BD Biosciences | Cat#: 555899 |
| BD Rhapsody™ Cartridge Kit | BD Biosciences | Cat#: 633733 |
| BD Rhapsody™ Cartridge Reagent Kit | BD Biosciences | Cat#: 633731 |
| BD™ Human Single-Cell Multiplexing Kit | BD Biosciences | Cat#: 633781 |
| BD Rhapsody™ WTA Amplification Kit | BD Biosciences | Cat#: 633801 |
| BD Rhapsody™ cDNA Kit | BD Biosciences | Cat#: 633773 |
| AMPure XP | Beckman Coulter | Cat#: A63880 |
| Qubit™ dsDNA HS Assay Kit | Invitrogen | Cat#: Q32854 |
| High Sensitivity D1000 Reagents | Agilent | Cat#: 5067-5585 |
| High Sensitivity D5000 Reagents | Agilent | Cat#: 5067-5593 |
| High Sensitivity D1000 ScreenTape | Agilent | Cat#: 5067-5584 |
| High Sensitivity D5000 ScreenTape | Agilent | Cat#: 5067-5588 |
| BD Pharmingen™ 7-AAD | BD Biosciences | Cat#: 559925 |
| Calcein AM | Invitrogen | Cat#: C1430 |
| Draq7 | BD Biosciences | Cat#: 564904 |
| Opal 7-Color Automated | Akoya Biosciences | Cat#: NEL821001KT |
| Immunohistochemistry Kit |  |  |
| BOND Epitope Retrieval 1 | Leica Biosystems | Cat#: AR9961 |
| BOND Epitope Retrieval 2 | Leica Biosystems | Cat#: AR9640 |
| BOND Dewax Solution | Leica Biosystems | Cat#: AR9222 |
| BOND Wash Solution 10x | Leica Biosystems | Cat#: AR9590 |
| Spectral DAPI | Akoya Biosciences | Cat#: FP1490 |
| Prolong Diamond Antifade | Thermo Fisher | Cat#: P36961 |
| BOND Research Detection System | Leica Biosystems | Cat#: DS9455 |
| BOND Titration Kit | Leica Biosystems | Cat#: OPT9049 |
| Antibodies used for flow cytometry |  |  |
| CD123 (clone 7G3) | BD Biosciences | Cat#: 563405;<br>RRID: AB_2738185 |
| CD14 (clone MφP9) | BD Biosciences | Cat#: 563372;<br>RRID: AB_2744290 |
| CD15 (clone HI98) | BD Biosciences | Cat#: 563838;<br>RRID: AB_2738444 |
| CD16 (clone eBioCB16) | Thermo Fisher | Cat#: 47-0168-42;<br>RRID: AB_11220086 |
| CD161 (clone DX12) | BD Biosciences | Cat#: 556081;<br>RRID: AB_396348 |
| CD181 (clone 8F1) | Miltenyi Biotec | Cat#: 130-105-353;<br>RRID: AB_2655708 |
| CD19 (clone HI98) | BD Biosciences | Cat#: 564977;<br>RRID: AB_2744308 |
| CD193 (clone 5E8) | BD Biosciences | Cat#: 562571;<br>RRID: AB_2737660 |
| CD28 (clone CD28.2) | BD Biosciences | Cat#: 751477;<br>RRID: AB_2875473 |
| CD3 (clone UCHT) | BD Biosciences | Cat#: 612940;<br>RRID: AB_2870222 |
| CD31 (clone WM59) | Biolegend | Cat#: 303122;<br>RRID: AB_2562149 |
| CD326 (clone 9C4) | Biolegend | Cat#: 324222;<br>RRID: AB_2561506 |
| CD34 (clone 8G12) | BD Biosciences | Cat#: 345801;<br>RRID: AB_2868825 |
| CD38 (clone HIT2) | BD Biosciences | Cat#: 562444;<br>RRID: AB_11151894 |
| CD4 (clone SK3) | BD Biosciences | Cat#: 612748;<br>RRID: AB_2870079 |
| CD45 (clone HI30) | BD Biosciences | Cat#: 564914;<br>RRID: AB_2744401 |
| CD49b (clone REA188) | Miltenyi Biotec | Cat#: 130-100-331;<br>RRID: AB_2658461 |
| CD54 (clone 84H10) | Beckman Coulter | Cat#: IM0726;<br>RRID: AB_130995 |
| CD56 (clone NCAM16.2) | BD Biosciences | Cat#: 563554;<br>RRID: AB_2687886 |
| CD62L (clone DREG-56) | BD Biosciences | Cat#: 555544;<br>RRID: AB_395928 |
| CD8 (clone RPA-T8) | BD Biosciences | Cat#: 612914;<br>RRID: AB_2870199 |
| CD83 (clone HB15e) | Thermo Fisher Scientific | Cat#: MHCD8301;<br>RRID: AB_10392703 |
| CD90 (clone 5E10) | Biolegend | Cat#: 328114;<br>RRID: AB_893431 |
| HLA-DR (clone G46-6) | BD Biosciences | Cat#: 556642;<br>RRID: AB_396508 |
| LOX1 (clone 15C4) | Biolegend | Cat#: 358604;<br>RRID: AB_2562181 |
| TCRgd (clone 5A6.E9) | Thermo Fisher Scientific | Cat#: MHGD06;<br>RRID: AB_10375477 |
| Antibodies used for multiplex IF staining |  |  |
| CD16 (clone EPR22409-124) | Abcam | Cat#: ab246222;<br>RRID: AB_2910166 |
| CXCR2 (clone EPR22301-103) | Abcam | Cat#: ab225732;<br>RRID: AB_2910167 |
| CXCR4 (clone EPUMBR3) | Abcam | Cat#: ab181020;<br>RRID: AB_2910168 |
| Cytokeratin (clone AE1/AE3) | Agilent | Cat#: M351501-2;<br>RRID: AB_2132885 |
| Cytokeratin (clone C-11) | Abcam | Cat#: ab7753;<br>RRID: AB_306047 |
| LOX-1/OLR1 (clone polyclonal) | Sigma-Aldrich | Cat#: HPA050798;<br>RRID: AB_2756318 |
| Deposited data |  |  |
| BD Rhapsody dataset (demultiplexed UMI counts) | This study | https://doi.org/10.5281/zenodo.6411868 |
| Processed input data | This study | https://doi.org/10.5281/zenodo.6411868 |
| Results, including intermediate results, core and extended atlas in h5ad format, and scArches model | This study | https://doi.org/10.5281/zenodo.6411868 |
| Core and extended atlas as cell-x-gene instance | This study | https://luca.icbi.at |
| Adams_Kaminski_2020<br>scRNA-seq dataset (processed) | GEO | GSE136831 |
| Chen_Zhang2020 scRNA-seq<br>dataset (fastq) | SRA | PRJNA634159 |
| Goveia_Carmeliet_2020<br>(processed) | (Goveia et al., 2020) | https://endotheliomics.shinyapps.io/lung_ectax/ |
| Guo_Zhang_2018 (fastq) | EGA | EGAS00001002430 |
| Habermann_Kropski_2020<br>(processed) | GEO | GSE135893 |
| He_Fan_2021 (fastq) | NGDC GSA | CRA001963 |
| Kim_Lee_2020 (processed) | GEO | GSE131907 |
| Lambrechts_Thienpont_2018<br>(fastq) | ArrayExpress | E-MTAB-6149, E-MTAB-6653 |
| Laughney_Massague_2020<br>(processed) | GEO | GSE123904 |
| Madissoon_Meyer_2020<br>(processed) | (Madissoon et al., 2019) | https://www.tissuestabilitycellatlas.org/ |
| Maier_Merad_2020<br>(processed) | (Maier et al., 2020) | https://github.com/effiken/Maier_et_al_nature_2020 |
| Maynard_Bivona_2020<br>(fastq) | SRA | PRJNA591860 |
| Mayr_Schiller_2020<br>(processed) | (Mayr et al., 2020) | https://github.com/theislab/2020_Mayr |
| Reyfman_Misharin_2018<br>(processed) | GEO | GSE122960 |
| Travaglini_Krasnow_2020<br>(processed) | (Travaglini et al., 2020) | https://www.synapse.org/#!Synapse:syn21560406 |
| Vieira_Teichmann_2019<br>(processed) | GEO | GSE130148 |
| Wu_Zhou_2021 (processed) | GEO | GSE148071 |
| Zillionis_Klein_2019<br>(processed) | GEO | GSE127465 |
| Leader_Merad_2021<br>(processed) | GEO | GSE154826 |
| TCGA data RNA-seq and<br>mutation data | GDC | https://portal.gdc.cancer.gov/ |
| TCGA response data | (Moiso, 2021) |  |
| TCGA survival data | (Liu et al., 2018) |  |
| OAK and POPLAR cohorts; clinical and RNA-seq data | EGA | EGAS00001005013 |
| LCAM signature | (Leader et al., 2021) | https://github.com/effiken/Leader_et_al/blob/3eb043acdea7b3dea45a2325ee1035060d9964af/scripts/get_LCAM_scores.R |
| Cytosig signatures | (Jiang et al., 2021) | https://github.com/data2intelligence/CytoSig/ |
| CellPhoneDB | (Efremova et al., 2020; Turei et al., 2021) | Downloaded from https://omnipathdb.org/interactions/?fields=sources,references&genesymbols=1&databases=CellPhoneDB on 2022-04-06 |
| Software and algorithms |  |  |
| Seven Bridges - BD Rhapsody™ WTA Analysis Pipeline | Seven Bridges Genomics | v1.7.1 |
| FlowJo™ | BD Biosciences | v10.7 |
| Mantra Snap | Akoya Biosciences | v1.0.4 |
| inForm Tissue Analysis | Akoya Biosciences | v2.4.10 |
| Cellranger v5.0.0 | 10x Genomics | https://support.10xgenomics.com/single-cell-gene-expression/software/pipelines/latest/what-is-cell-ranger |
| nf-core RNA-seq pipeline v3.0 | (Patel et al., 2022) | https://github.com/nf-core/rnaseq |
| Nextflow v21.10.0 | (Di Tommaso et al., 2017) | https://nextflow.io |
| Singularity/Apptainer v3.7.0-1.el7 | (Kurtzer et al., 2017) | https://apptainer.org/ |
| Nextflow workflow to reproduce this study | This study | https://github.com/icbi-lab/luca |
| Software packages used for scRNA-seq analysis are packaged as singularity containers and available on zenodo | This study | https://doi.org/10.5281/zenodo.6411868 |
| Compute hardware |  |  |
| Compute node CPU | Intel | Xeon(R) CPU E5-2699A v4 (2x) |
| GPU node GPU | Nvidia | Quadro RTX 8000 |
| GPU node CPU | AMD | EPYC 7352 24-Core (2x) |

## STAR METHODS

### Resource availability

#### Lead contact

Further information and requests for resources and reagents should be directed to and will be fulfilled by the lead contact, Zlatko Trajanoski

#### Materials availability

The study did not generate new unique reagents.

#### Data and code availability

- Processed scRNA-seq data from this study has been deposited on Zenodo as listed in the key resources table. Raw data is not made available due to privacy concerns.
- Processed scRNA-seq data from other studies has been deposited on Zenodo. The original study identifiers are listed in the key resource table.
- Final and intermediate results of the computational analysis are made available on Zenodo.
- All code to reproduce this study is wrapped into a nextflow workflow and publicly available on Github. All software dependencies are made available as singularity containers. Some of the algorithms employed (scVI, scANVI, UMAP) involve stochastic processes that require specific hardware for exact reproducibility (see key resource table).

#### Human subjects

Samples of NSCLC tumor tissues and matched adjacent normal lung tissues (more than 5 cm distance to the tumor) were obtained from surgical specimens of patients undergoing resection at the Department of Visceral, Transplant and Thoracic Surgery (VTT), Medical University Innsbruck, Austria, and in collaboration with the INNPATH GmbH, Innsbruck, Austria, after obtaining informed consent in accordance with a protocol reviewed and approved by the Institutional Review Board at the Medical University Innsbruck, Austria (study code: AN214-0293 342/4.5). Demographic details are provided in Table S5.

#### Sample preparation of NSCLC tissue and corresponding adjacent normal lung tissue

Surgically resected NSCLC tumor tissues and adjacent normal tissues were minced into small pieces (< 1 mm) on ice and enzymatically digested with agitation for 30 minutes at 37°C using the BD TuDoR™ dissociation reagent (BD Biosciences). The obtained single-cell solution was sieved through a 70 µM cell strainer (Corning) and red blood cells were removed using the BD Pharm Lyse™ lysing solution (BD Biosciences). Cells were counted and viability assessed with the BD Rhapsody scRNA-seq platform (BD Biosciences) using Calcein-AM (Invitrogen) and Draq7 (BD Biosciences).

#### BD Rhapsody scRNA-seq library preparation and sequencing

Freshly isolated single-cells were immediately processed with the BD Rhapsody scRNA-seq platform (BD Biosciences). The BD Single-Cell Multiplexing Kit (BD Biosciences) was used to combine and load two samples (tumor tissue and normal adjacent tissue) onto a single BD Rhapsody™ cartridge (BD Biosciences). Sample-tag staining was performed according to the manufacturer’s protocol (sample-tag staining at room temperature for 20 minutes and washing by centrifugation at 400 g for 5 minutes). Single-cell isolation in microwells (cell load: 20 minutes incubation at room temperature) with subsequent cell-lysis and capturing of poly-adenylated mRNA molecules with barcoded, magnetic captured-beads was performed according to the manufacturer’s instructions. Beads were magnetically retrieved from the microwells, pooled into a single tube before reverse transcription. Unique molecular identifiers (UMIs) were added to the cDNA molecules during cDNA synthesis. Whole transcriptome amplification (WTA) and sample-tag sequencing libraries were generated according to the BD Rhapsody single-cell whole-transcriptome amplification workflow. The quantity and quality of the sequencing libraries was analyzed with the Qubit dsDNA HS (High Sensitivity) assay kit (Invitrogen) and the 4200 TapeStation (Agilent) system. Libraries were sequenced on the Novaseq 6000 system (Illumina) targeting a sequencing depth of 50.000 reads/cell.

#### Flow cytometry

Cells isolated from surgically resected NSCLC tumor tissues and adjacent normal tissues were stained with a backbone cocktail of 12 antibodies which, was complemented either with an additional 8 antibodies to define all cell populations, or several mixtures of up to three antibodies for a detailed characterization of neutrophils at pre-titrated concentrations (Table S6). After washing and addition of 5 µl 7-AAD, the cells were measured on a FACSymphony A5 flow cytometer (BD Biosciences). Data were analyzed using FlowJo v10.7 software. For details of the gating strategy see Figure S7.

#### Multiplex immunofluorescence (IF) staining and analysis

NSCLC tumor and tumor-adjacent tissue samples were fixed in 4% formalin for 6-72 hours and embedded in paraffin. Four-micrometer sections were used for the immunofluorescence staining. Immunofluorescence staining on Formalin-fixed paraffin-embedded (FFPE) tissue was performed using antibodies against cytokeratin (Agilent, dilution 1:500; Abcam, dilution 1:1000), CXCR2 (Abcam, dilution 1:500) and LOX-1 (Sigma-Aldrich, dilution 1:200). The staining procedure was performed using an automated staining system (BOND-RX, Leica Biosystems). All markers were sequentially applied and paired with respective Opal fluorophores (Table S6). To visualize cell nuclei, the tissue was stained with 4‘,6-dia-midino-2-phenylindole (spectral DAPI, Akoya Biosciences). Stained slides were scanned using Mantra 2 Quantitative Pathology Workstation (Akoya Biosciences) and representative images from each tissue were acquired with the Mantra Snap software v1.0.4. Spectral unmixing, multispectral image analysis and cell phenotyping was carried out using the inForm Tissue Analysis Software v2.4.10 (Akoya Biosciences).

### Quantification and statistical analysis

#### Preprocessing and quality control of sequencing data

We distinguish between studies (i.e. a scientific publication) and datasets (i.e. scRNA-seq samples that were generated using the same sample preparation and the same experimental platform). Each study may contain one or multiple datasets. Demultiplexed FASTQ files of the UKIM-V datasets were merged and processed using the Seven Bridges Genomics cloud server with the BD Rhapsody WTA Analysis Pipeline. Samples from the studies Chen_Zhang_2020, Guo_Zhang_2018, He_Fan_2021, Lam-brechts_Thienpont_2018 and Maynard_Bivona_2020 were obtained as raw fastq files from the identifiers specified in the key resource table. Smart-seq2 data were processed using the nf-core RNA-seq pipeline (Ewels et al., 2020; Patel et al., 2022) with the GRCh38 reference genome and GENCODE v33 annotations. 10x datasets were processed with cellranger v5.0.0 (10x Genomics) and the GRCh38-2020-A reference database as provided by 10x Genomics. All other datasets were obtained as count tables from their respective identifiers. All datasets were loaded into AnnData containers (Virshup et al., 2021) with consistent structure. Quality control was performed with scanpy (Wolf et al., 2018) by thresholding the number of detected genes, counts and the fraction of mitochondrial reads. Thresholds were determined per dataset by visual inspection of the distributions and are listed in Table S7.

#### Integration of scRNA-seq datasets

Individual datasets were merged into a single AnnData object. Since genome annotations partly differed between the datasets, we re-mapped gene identifiers on the latest version of HGNC gene symbols using the https://mygene.info API (Xin et al., 2016). In case of duplicate gene symbols, the one with the maximum read count was retained. If gene symbols were missing from a dataset, the values were filled with zeros. Gene symbols that were missing in more than 5 datasets (25%) were excluded altogether.

We integrated the datasets using the scANVI algorithm (Gayoso et al., 2022; Xu et al., 2021), as it has been demonstrated to be one of the top-performing methods for atlas-level integration and to scale to >1M cells (Luecken et al., 2022). Since scANVI requires cell-type annotations for at least one of the input datasets, we manually annotated two “seed” datasets based on unsupervised clustering as described below. We chose Lambrechts_Thienpont_2018_6653 and Maynard_Bivona_2020 as seed datasets as they were not experimentally enriched for specific cell-types and were sequenced on two platforms with very different characteristics (10x and Smart-seq2). Raw counts were used as input for scANVI. The Smart-seq2 counts were scaled by the gene length as recommended on the scvi-tools website. The scANVI model was initialized with a pre-trained scVI model (Lopez et al., 2018), as recommended in the scvi-tools tutorial. The scVI model was trained on the 6000 most highly variable genes as determined with scanpy’s (Wolf et al., 2018) *pp.highly_variable_genes* with parameters *flavor=”seurat_v3”* and *batch_key=”dataset”*. Each sample was considered as an individual batch for both scVI and scANVI. Other than that the algorithms were run with default parameters.

#### Doublet-detection

For droplet-based scRNA-seq datasets we ran the SOLO algorithm (Bernstein et al., 2020) to computationally detect multiplets. We chose SOLO over other doublet detection methods as it is readily integrated into scvi-tools (Gayoso et al., 2022), and was found to be one of the top-performing methods in an independent benchmark (Xi and Li, 2021). We used the SOLO implementation from scvi-tools and initialized SOLO with a pre-trained scVI model.

#### Unsupervised clustering and cell-type annotation

We computed UMAP embeddings (Becht et al., 2018) and unsupervised Leiden-clustering (Traag et al., 2019) with scanpy (Wolf et al., 2018), based on a cell-cell neighborhood graph derived from scANVI latent space. Coarse, lineage-specific clusters were iteratively sub-clustered to identify cell-types at a more fine-grained resolution. Cell type clusters were annotated based on previously reported marker genes (Madissoon et al., 2019; Schupp et al., 2021; Sikkema et al., 2022) (Figure S1A).

#### Integrating additional datasets

Two datasets, Leader_Merad_2021 and UKIM-V-2, were added after the completion and annotation of the core atlas. The datasets (“query”) were projected onto the atlas (“reference”) using scArches (Lotfollahi et al., 2022) as implemented in scvi-tools (Gayoso et al., 2022). scVI and scANVI models were re-trained on the fully annotated, doublet-filtered core atlas, with the parameters recommended for scArches: *use_layer_norm=”both”, use_batch_norm=”none”, encode_covariates=True, drop-out_rate=0.2, and n_layers=2.* Gene-symbols of the query datasets were re-mapped as described above and missing gene symbols filled with zeros. For each query dataset, scArches yielded an embedding in the same latent space as the core atlas. Based on the joint latent space, a neighborhood graph and UMAP embedding were computed for the “extended” atlas. Cell-types were annotated automatically, based on a majority vote of nearest neighbors. To this end, let 𝐶 be the pairwise weighted connectivity matrix of the scanpy neighborhood graph computed on the scArches embedding. Then, the transitive connectivity matrix 𝐶′ (i.e., including connections to neighbors of neighbors) is defined as 𝐶′ = 𝐶 · 𝐶 where the dot operator refers to the matrix product. Let further 𝑄 be the set of all query cells, 𝑅 the set of all reference cells, and 𝑇 the set of all cell-types. Then, for every cell 𝑞 ∈ 𝑄 the cell-type is determined as

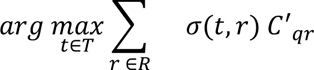

where the indicator function 𝜎(𝑡, 𝑟) is 1 if cell *r* is of type *t* and 0 otherwise. The transitive connectivity matrix 𝐶′ was chosen over 𝐶 to increase robustness by increasing the number of neighbors, and to ensure that every cell from the query has connection to a cell in the reference.

#### Comparing cell-type abundances

Since comparing cell-type fractions between groups is challenging due to different characteristics of the datasets and the inherent compositional nature of cell-type fractions, we applied the scCODA (Buttner et al., 2021) model, which addresses this issue. We were interested in the differences between conditions (LUAD vs LUSC). To this end, we ran the scCODA model with the formula *∼ condition + tumor_stage + dataset* with 500,000 iterations using “Tumor cells” as the reference cell-type, where *tumor_stage* is a binary vector classifying datasets into early (stages I-II) and advanced (stages III-IV), and dataset is a categorical vector encoding the different datasets. For the comparison, we excluded the Guo_Zhang_2018 dataset, which only contains T cells. The final result shows *credible effects* with a false-discovery-rate (FDR) of 0.1.

#### Patient stratification

We stratified patients into immune phenotypes based on cell-type fractions. We selected all patients with primary tumor samples and excluded the Guo_Zhang_2018 dataset, because it contains only T cells. Cell-type fractions of primary tumor samples were loaded into a patient × cell-type AnnData container. Dataset-specific batch-effects were removed using a linear model as implemented in *scanpy.pp.regress_out*. Patients were clustered using graph-based Leiden clustering with the “correlation” distance metric for computing the neighborhood graph. Patient clusters were labeled according to their predominant cell-types. In addition to the histological subtypes based on the annotation of the original datasets, we annotated tumor types based on the single-cell data according to the most abundant tumor cell-type (LUAD, LUSC, LUAD NE, LUAD EMT and LUAD dedifferentiated).

#### RNA Velocity analysis

We performed RNA-velocity analysis on the UKIM-V dataset using velocyto.py (La Manno et al., 2018) and scvelo (Bergen et al., 2020). BAM files as generated by the BD Rhapsody WTA analysis pipeline were preprocessed with samtools (Danecek et al., 2021) to make them compatible with velocyto.py (see preprocessing/bd_rhapsody/velocyto.nf in our git repository for more details). Loom files generated by velocyto.py were loaded into scvelo to estimate and visualize RNA velocities according to the scvelo tutorial. Partition-based graph abstraction (PAGA, (Wolf et al., 2019)) was computed based on the RNA velocity graph, using neutrophil subclusters as grouping variable and the option *minium_spanning_tree=False*. The result was visualized as a graph showing the transition confidences as directed edges.

#### Differential gene expression testing

We used DESeq2 (Love et al., 2014) on pseudo-bulk samples for differential expression testing which has been demonstrated to perform well and properly correct for false discoveries (Squair et al., 2021). For each cell-type and patient, we summed up transcript counts for each gene. Pseudo-bulk samples consisting of fewer than 10 cells were discarded. Primary tumor samples from LUAD *vs.* LUSC were compared using the linear model *∼ condition + dataset*. Primary tumor samples from the patient subtypes, M, B and T were independently compared to the “deserted” subtype using the linear model *∼ group + dataset.* P-values were adjusted for multiple hypothesis testing with independent hypothesis weighting (IHW) (Ignatiadis et al., 2016).

#### Pathway, transcription factor and cytokine signaling signatures

We performed pathway, transcription factor, and cytokine signaling analysis on primary tumor samples with PROGENy (Holland et al., 2020; Schubert et al., 2018), DoROthEA (Garcia-Alonso et al., 2019; Holland et al., 2020) and CytoSig (Jiang et al., 2021), respectively. Scores were computed using the *dorothea-py* and *progeny-py* packages. The top 1,000 target genes of the progeny model were used, as recommended for single-cell data. For dorothea, only regulons of the highest confidence levels “A” and “B” were used. The cytosig signature matrix was obtained from the *data2intelligence/CytoSig* GitHub repository and used with the scoring function implemented in the *progeny-py* package. The methods were run with the options *num_perm=0, center=True, norm=True scale=True,* and *min_size=5.* No permutations were used, as we perform statistics in a separate step at the level of biological replicates. Pathway-, transcription factor-, and cytosig scores were then compared between *condition* (LUAD vs. LUSC) and patient *group* (T vs. B vs M vs deserted) using an ordinary least-squares (OLS) linear model, as implemented in the statsmodels package (Seabold and Perktold, 2010). Scores were aggregated into pseudobulk samples by computing the mean of each variable for each patient and cell-type. Samples consisting of less than 10 cells were discarded. For each variable, we fitted a model with the formulas *∼ condition + dataset + tumor_stage* or *∼ group + dataset + condition + tumor_stage*, respectively. Coefficients were obtained from the linear model and p-values calculated with the f-test. P-values were adjusted for multiple testing with the Benjamini-Hochberg procedure.

#### Cellphonedb analysis

We used the cellphonedb (CPDB) database (Efremova et al., 2020) as obtained from omnipathdb (Turei et al., 2021) to investigate differences in cell-to-cell communication in primary tumor samples. The original CPDB algorithm performs statistical comparisons based on a permutation test which is designed to find differences between cell-types. For our study, on the other hand, we were interested in differences between conditions, using patients as biological replicates. Therefore, we followed an approach similar to the *degs_analysis* mode recently added to CPDB v3 (Garcia-Alonso et al., 2021): For each cell-type of interest, we considered the list of significantly differentially expressed signaling molecules in CPDB (ligands or receptors, for outgoing and incoming interactions, respectively). For each of those differentially expressed signaling molecules and for each cell-type, we determined interaction partners that are potentially affected by that change, as those that are expressed in at least 10% of the cells in a certain cell-type. Differentially expressed signaling molecules were determined with DESeq2 as described above (for LUAD vs. LUSC and the comparison between patient subtypes), or ordinary least squares regression on log-normalized pseudo-bulk expression values with sum-to-zero coding (for a multi-group comparison between neutrophil subclusters). The fraction of cells expressing a signaling molecule was computed as the mean of fractions per patient, to avoid biases due to different cell-counts per patient

#### SCISSOR analysis

We used SCISSOR (Sun et al., 2022) to associate phenotypic data from bulk RNA-seq experiments with our single-cell data. TCGA mutation and gene expression data was obtained from the GDC portal, survival data from (Liu et al., 2018). SCISSOR was run on each patient individually according to the scissor tutorial using mutation data (logistic regression) and overall survival (cox-regression) as dependent variables. A grid search for the alpha-parameter was performed in 2^-i/2^ with 𝑖 ∈ [24, 23, . . ., 2] and a cutoff parameter of 0.3. 43 of 309 samples with low overall cell count failed during SCISSOR’s Seurat-preprocessing step and were excluded from the subsequent analysis. For each patient and cell-type, we computed the fraction of *scissor+ cells* (i.e. positively associated with a mutation or worse survival), *scissor- cells* (i.e. negatively associated), and *neutral cells* and added a pseudo-count of 0.01. A sample was excluded from a cell-type if it contributed ≤10 cells. For each cell-type, we computed the log2-ratio of scissor+ and scissor-cells as the mean fraction of scissor+ cells *vs.* the mean fraction of scissor-cells. Significant differences were determined by comparing the fractions of scissor+ and scissor-cells with a paired wilcoxon test with *zero_method=”zsplit”* as implemented in the scipy package. P-values were Benjamini-Hochberg-adjusted and considered significant at an FDR < 0.01.

#### Copy number inference

We used SCEVAN (De Falco et al., 2021) to infer copy number aberrations (CNA) from single-cell gene transcriptomics data. SCEVAN was run individually for each patient using raw counts as input and with disabled subclone calling. The resulting CNA matrices were grouped into the respective immune phenotypes (Immune deserted, B infiltrated, M infiltrated and T infiltrated). For each patient and segment, the mean log2(CNR) was calculated across all cells, where *CNR* refers to the copy number ratio. Significant CNR differences between the immune phenotype groups were determined by fighting a linear regression model using robust Heteroscedasticity Consistent version 3 (*HC3*) standard errors with the formula *∼ group + dataset + condition + tumor_stage*. The results were filtered for |log2(CNR)| > 0.01 and FDR < 0.1 and circos plots were produced using the R circlize library. We calculated intratumoral heterogeneity scores for CNA (𝐻_CNA_) and gene expression (𝐻_GEX_) (Wu et al., 2021) based on SCEVAN’s CNA matrix as implemented in *infercnvpy* (https://github.com/icbi-lab/infercnvpy). To assess the association of ITHGEX/ITHCNA with tumor stage, immune phenotype and tumor type, we fitted an ordinary least squares linear model with the formula *∼ immune_phenotype + tumor_type + tumor_stage + dataset + number_of_tumor_cells*.

#### TCGA copy number analysis

We used k-means clustering with the cell type fractions available from TCIA (https://tcia.at, (Charoen-tong et al., 2017)) to partition LUAD and LUSC patient samples from TCGA into two groups, immune infiltrated and immune deserted. For each sample we retrieved the masked copy number segment data from GDC via the R Bioconductor package GenomicsDataCommons and assigned the “Segment_Mean” log2(CNR) to each gene overlapping with a given segment. Using the Wilcoxon rank sum test we determined significant copy number ratio differences between protein coding genes in the immune infiltrated and the immune deserted subtype and plotted them using the R circlize library (p.adj < 0.0001, and |log2(CNR)| > 0.1). We further filtered the genes, keeping only those with a |log2(CNR)| > 0.3.

#### TRN clusters

For an unbiased discovery of TRN subtypes, we performed unsupervised clustering of all cells annotated as neutrophils. The neighborhood graph was computed with *scanpy.pp.neighbors* with *n_neighbors=20* based on the scANVI latent space. Clusters were determined with *scanpy.tl.leiden* with *resolution=0.5*. Two subclusters dominated by cells from normal adjacent tissue were labeled normal-associated neutrophils (NAN) 1 and 2, whereas four subclusters of cells from primary tumor samples were labeled tumor-associated neutrophils (TAN) 1, 2, 3 and 4.

#### TRN signatures

Gene signatures for TRN and TRN clusters were determined based on fold-change (FC), specific fold-change (sFC), and area under the receiver operator characteristics curve (AUROC), applying an approach previously used to find cell-type-specific marker genes (Becht et al., 2016). We have previously shown the resulting gene signatures to be highly specific for their respective cell-types (Sturm et al., 2019). To avoid marker genes being biased towards samples contributing more cells than on average we aggregated single cells to pseudo-bulk samples (Squair et al., 2021) by patient before deriving marker genes. For each set of marker genes derived, pseudo-bulk samples were generated by summing up raw counts for each patient and cell-type of interest.The resulting samples were normalized to counts per million (CPM) and log2-transformed with *scanpy.pp.log1p(adata, base=2)*. Pseudo-bulk samples consisting of fewer than 10 cells were discarded. For each gene and cell-type, FC, sFC were computed as described in (Becht et al., 2016). AUROC was computed using *roc_auc_score* as implemented in scikitlearn. We defined signatures to: (1) discriminate TANs from NANs, (2) discriminate the 6 neutrophil subclusters from each other, and (3) discriminate neutrophils from other cell-types. For the TRN signature, we applied cutoffs of FC > 2, sFC > 1.5 and AUROC > 0.97 for finding cell-type-specific marker genes for cell-type estimation from bulk RNA-seq samples as previously used (Becht et al., 2016), which resulted in 36 highly-specific marker genes (Figure 6H). To determine markers specific for TAN/NAN or neutrophil subclusters and used relaxed cutoffs of FC > 1.5, sFC > 1 and AUROC > 0.75 at the cost of reduced marker specificity with top markers reaching an AUROC of 0.86-0.99 depending on the neutrophil subtype. Signature scores in scRNA-seq data were computed using *scanpy.tl.score_genes*.

#### Signature scoring in bulk RNA-seq samples

Bulk RNA-seq primary tumor samples samples of TCGA LUAD and LUSC were retrieved as TPM from the GDC portal. TCGA drug response data was retrieved from (Moiso, 2021). Bulk RNA-seq samples from NSCLC patients treated with atezolizumab (anti-PD-L1) or docetaxel (chemotherapy) from the POPLAR (Fehrenbacher et al., 2016) and OAK (Rittmeyer et al., 2017) trials were retrieved using the accession numbers reported in (Patil et al., 2022). LCAM-consensus signature score was computed as described in (Leader et al., 2021) using the code provided in their GitHub repository. Similar to that approach, enrichment scores for our neutrophil signature were calculated as follows: For all signature genes, z-scores were computed across all samples from a dataset. The final signature score was defined as the mean of the z-scores of the signature genes for each sample. Associations of the TRN signature with response to immunotherapy or chemotherapy in the POPLAR and OAK datasets was tested using logistic regression in R with the formula *response ∼ signature_score + tumor_type + dataset,* where tumor_type represents LUAD and LUSC encoded as a binary vector.

#### Survival analysis

Survival analysis was performed using CoxPH-regression as implemented in the R package *survival.* Kaplan-Meyer plots were created using the R package *survminer*, showing the top 25% vs. bottom 25% of samples stratified by signature score. B cell fractions in TCGA samples were estimated using EPIC (Racle et al., 2017) as implemented in *immunedeconv*, as we have previously shown EPIC to be one of the best performing methods on B cells (Sturm et al., 2019). Cox-regression was performed on B cell fractions (TCGA data) with the formula *survival ∼ signature_score + ajcc_stage + age,* where ajcc_stage is a categorical vector with tumor stages I-IV. For neutrophil fractions (POPLAR+OAK data) the formula *survival ∼ signature_score + dataset + treatment* was used.

## SUPPLEMENTAL INFORMATION

### Supplemental Figure Legends

**Supplemental Figure 1.**

(A) Dotplot of cell type marker genes used for cell-type annotation.

(B) Fractions of cell types, sample origins and conditions per study (extended atlas).

(C) Relative cell type proportions by tissue origin in the core atlas. The depicted fractions are the average of all patients, independent of their cell-count.

(D) Marker genes for tumor cell classification.

(E) Cell type fractions in the extended atlas.

(F) Mean neutrophil fraction per sequencing platform across all datasets.

(G) Flow-cytometry of neutrophils (shown as percentage of leucocytes) in tumor tissue and patient-matched normal-adjacent tissue (n=63; Paired Wilcoxon test, **p<0.01). The horizontal line represents the median, whiskers extend to the inter-quartile range.

(H) Number of reads (Smart-seq2) or UMIs (other platforms) in epithelial cells by sequencing platform. The central line denotes the median, boxes represent the interquartile range (IQR) and whiskers extend to the most extreme values within 1.5 * IQR. Points outside 1.5 * IQR are shown as outliers.

(I) UMAP of the extended atlas, colored by the medium resolution cell-type annotation used for most data analyses.

**Supplemental Figure 2**

(A) Flow cytometry analysis of the T cell-to myeloid cell ratio (T/M ratio) in tumor tissue (T subtypen=4, M subtypen=2). The horizontal line represents the median, whiskers extend to the inter-quartile range.

(B) Fractions of immune phenotypes in histological subtypes.

(C) Differentially expressed transcription factors (TFs) of tumor cells across immune phenotypes. Heatmap colors indicate the deviation from the overall mean, independent of tumor histology and stage. White dots indicate significant interactions at different false-discovery-rate (FDR) thresholds. P-values have been calculated using a linear model f-test. Only TF signatures with an FDR < 0.1 in at least one patient subtypeare shown.

(D) Boxplots of intratumoral heterogeneity of copy number abberations (ITHCNA, upper panel) and gene expression (ITHGEX, lower panel) by histology, immune phenotypes and tumor stages. Each dot represents a patient with at least 50 tumor cells and is colored by the study of origin. The central line denotes the median, boxes represent the interquartile range (IQR) and whiskers extend to the most extreme values within 1.5 * IQR.

(E) Circos plot of significant CNAs for the the immune deserted and immune infiltrated tumors of the TCGA cohort. Highlighted are selected genes that match the CNAs in the single-cell NSCLC cohort. Significant genes are determined as having a log2(copy number ratio) > 0.1 and an FDR < 0.0001 (Wilcoxon test).

**Supplemental Figure 3.**

(A) Tumor-immune crosstalk in LUAD *vs.* LUSC (resembling Circos plot Figure3A). Upper panel: top 30 differentially expressed ligands in LUAD *vs.* LUSC (DESeq2 on pseudo-bulk, FDR < 0.01). Heatmap colors indicate log2 fold changes clipped at ±5, where blue indicates upgregulation in LUSC and red indicates upregulation in LUAD. Bottom panel: Respective receptors and the expression by cell type. Dot sizes and colors refers to the fraction of cells expressing the receptor and gene expression, respectively, averaged over all patients. Dots are only shown for receptors that are expressed in at least 10% of the respective cell-types.

**Supplemental Figure 4**

(A-B) Association of cellular composition and TP53 mutation in LUAD and LUSC derived from the TCGA reference dataset.

(C-D) Association of cellular composition and survival for LUSC and LUAD patients.

(E-F) Kaplan-Meyer plot of LUAD and LUSC patients with high (top 25%) and low (bottom 25%) B cell fractions of TCGA lung cancer patients as determined by deconvolution with EPIC. P-value has been determined using CoxPH-regression using tumor stage and age as covariates.

**Supplemental Figure 5**

(A) Neutrophil fraction in LUSC *vs.* LUAD (extended atlas). P-value derived using linear model f-test including dataset as a covariate.

(B) Expression of top 30 marker genes (AUROC > 0.75) for NANs and TANs. Every dot is the log2 fold change on a single patient. Bars show the average log2-fold change.

(C) Multiplex immunofluorescence co-staining of CXCR2 (red), LOX-1 (yellow) and pan-cytokeratin (blue) in NSCLC tumor tissue. Scale bar = 100 µm.

**Supplemental Figure 6.**

(A-B) TRN subclusters by the contributing datasets (A) and by patients (B). Data of 32 patients with each > 10 neutrophils.

(C) Selected TRN subluster genes.

(D) TRN signature gene expression across TRN subclusters.

(E) UMAP plot of atlas neutrophils colored by gene expression of selected marker genes for each TRN subcluster.

(F) Flow cytometry analysis demonstrating the correlation between HLA-DR expression and CD83, LOX-1, CD181, CD62L or CD16 expression, respectively. Representative analysis of neutrophils derived from NSCLC normal-adjacent tissue (blue) and tumor tissue (red) are shown.

(G) Kaplan-Meyer plot of LUSC patients form the POPLAR (Fehrenbacher *et al*., 2016) and OAK (Rittmeyer *et al*., 2017) cohorts treated with atezolizumab with high (top 25%) and low (bottom 25%) TRN signature score. P-value has been determined using CoxPH-regression.

(H) Kaplan-Meyer plot of LUAD patients form the POPLAR (Fehrenbacher *et al*., 2016) and OAK (Rittmeyer *et al*., 2017) cohorts treated with atezolizumab with high (top 25%) and low (bottom 25%) TRN signature score. P-value has been determined using CoxPH-regression.

(I) Correlations of the TRN with the LCAM-consensus (Leader *et al*., 2021) signature scores in bulk RNA-seq samplese from two NSCLC cohorts of the POPLAR (Fehrenbacher *et al*., 2016) and OAK (Rittmeyer *et al*., 2017) trials treated with atezolizumab colored by response to immunotherapy.

**Supplemental Figure 7.**

(A) Flow cytometry gating strategy to define cell populations from NSCLC tumor tissue and normal-adjacent tissue. In initial cleaning steps dead cells, debris and doublets were removed using 7-AAD staining and scatter characteristics. Leukocytes were defined by CD45 staining and sequentially gated into subtypes including neutrophils, monocytes, T cells and B cells. Non-CD45^+^ cells were gated into epithelial cells, endothelial cells and fibroblasts.

### Supplemental Tables

**Supplemental Table 1: Study characteristics.**

**Supplemental Table 2: Genes with significant CNA from the extended scRNA-seq atlas cohort. Supplemental Table 3: Genes with significant CNA from the TCGA cohort.**

**Supplemental Table 4: An overlap of genes with significant CNA from the extended scRNA-seq atlas cohort with genes with significant CNA from the TCGA cohort.**

**Supplemental Table 5: Patient metadata.**

**Supplemental Table 6: Dataset quality control thresholds. Supplemental Table 7: Immunofluorescence and FACS.**

