## Supplementary figures and images for "High-resolution single-cell atlas reveals diversity and plasticity of tissue-resident neutrophils in non-small cell lung cancer"

### Supplementary Figures

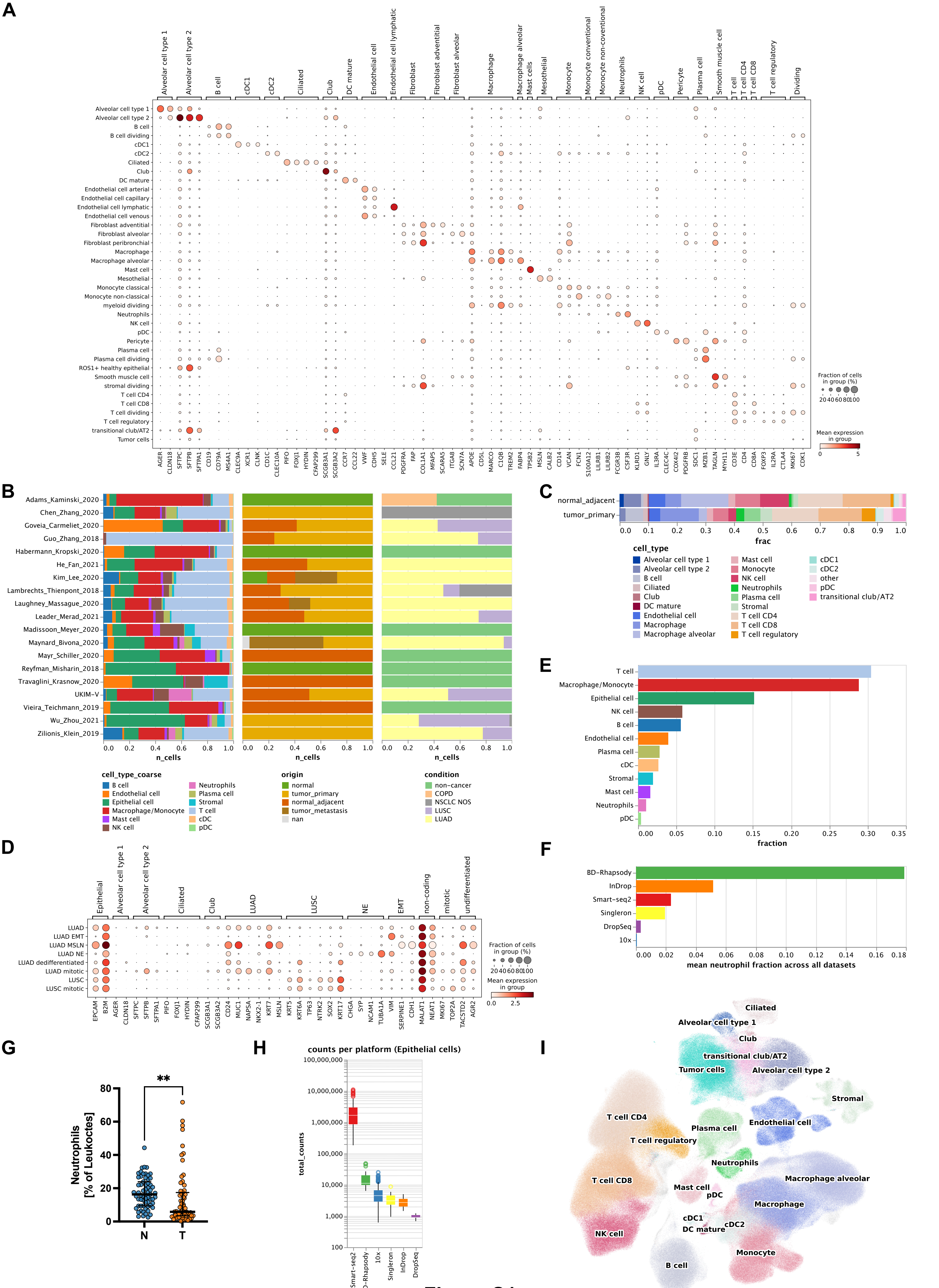

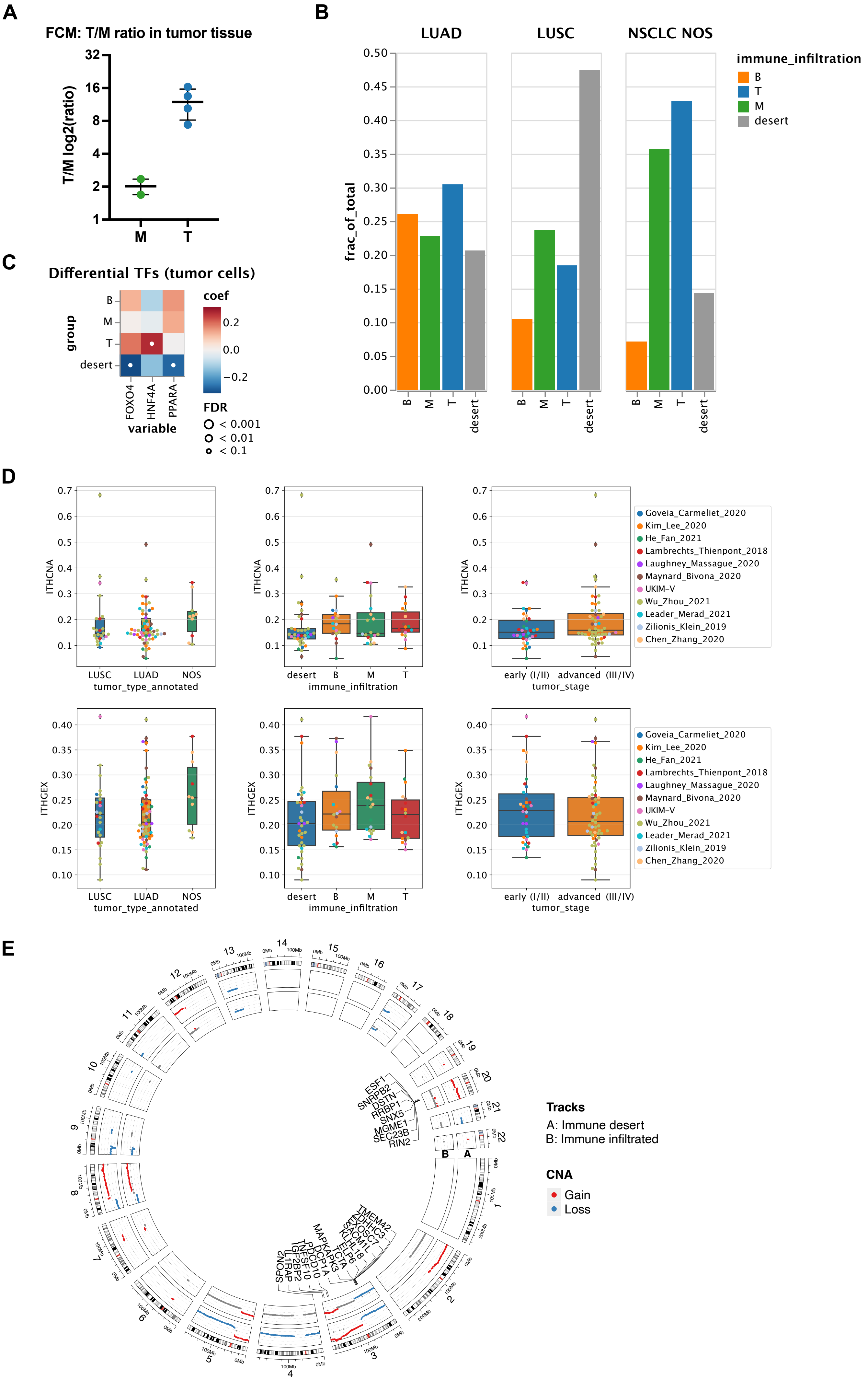

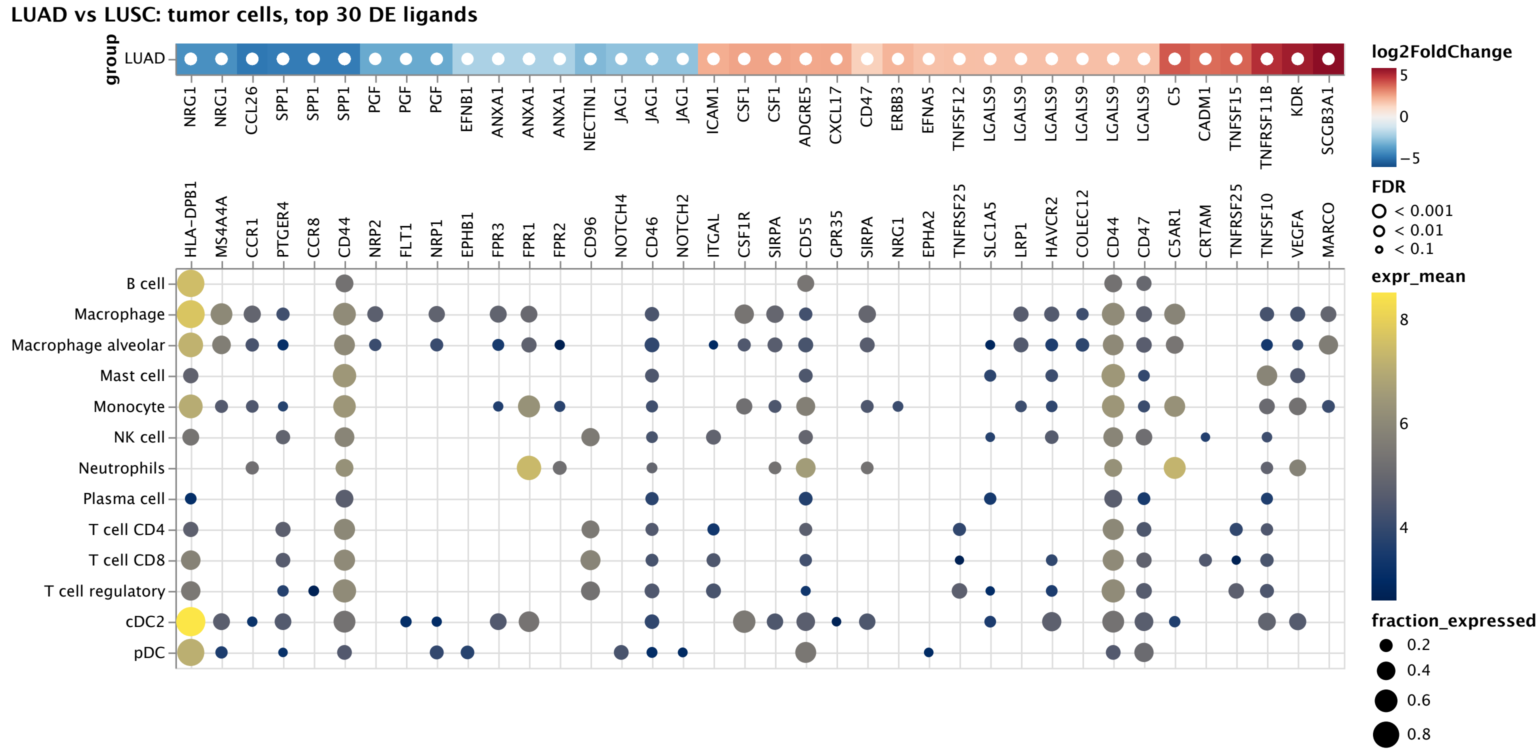

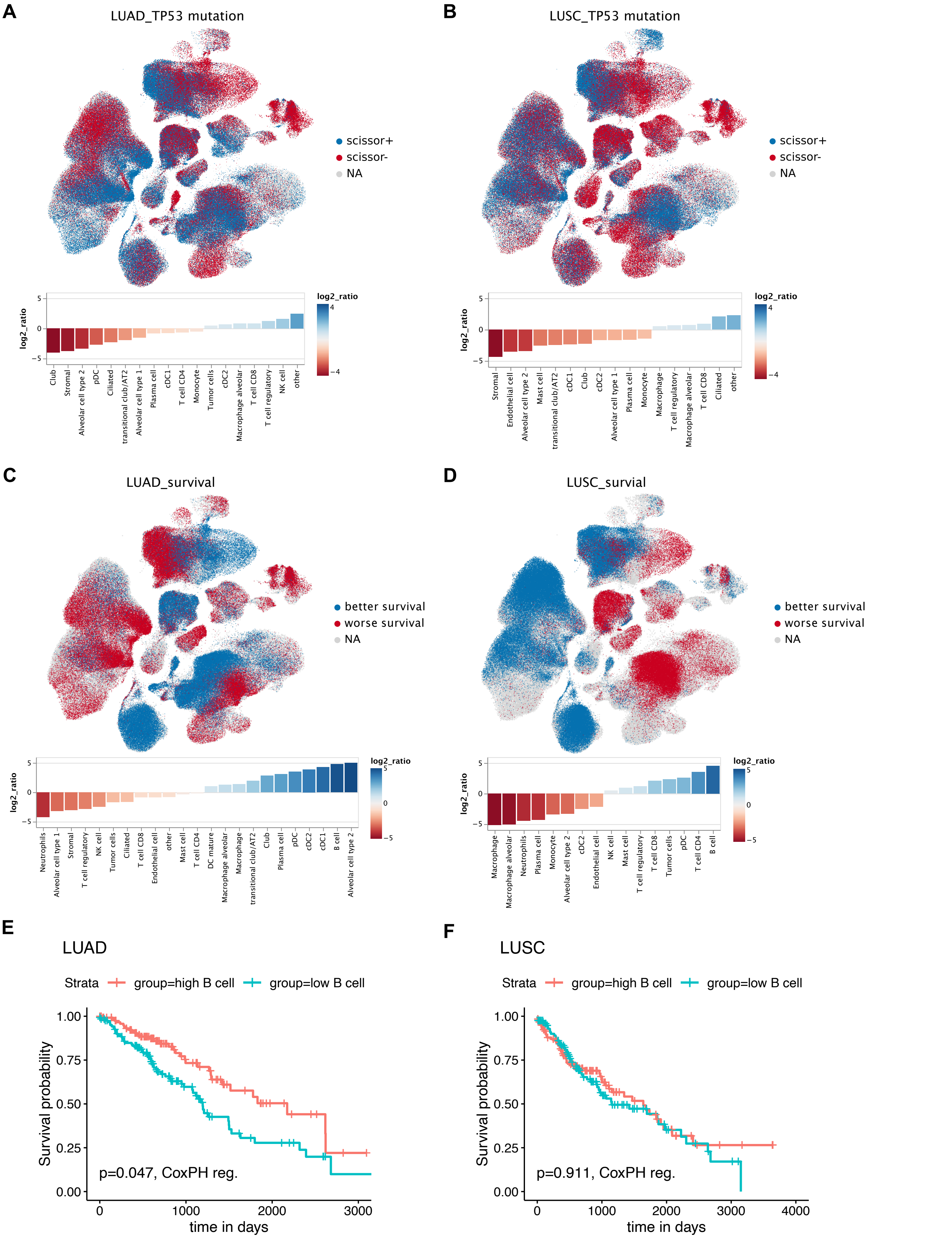

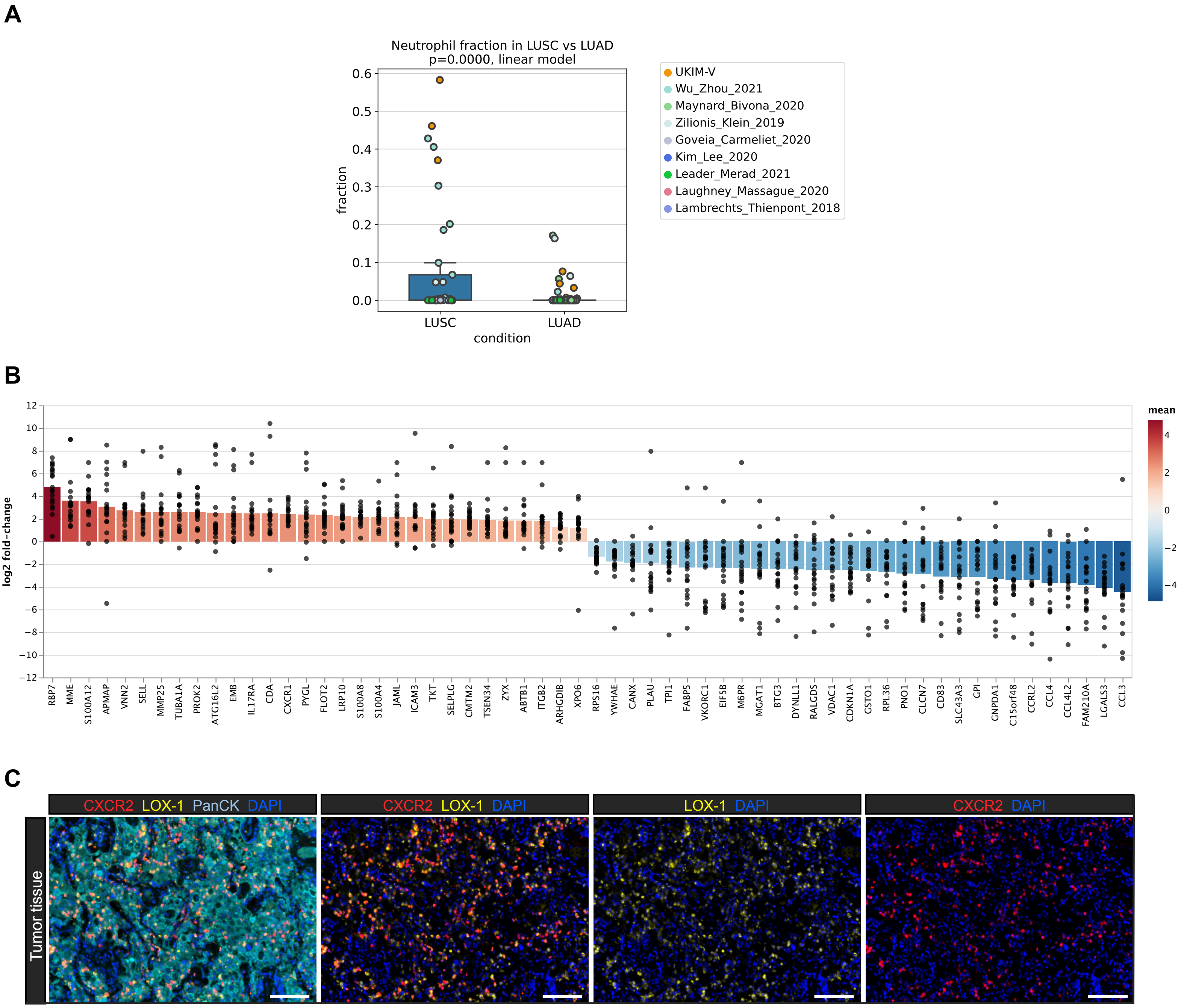

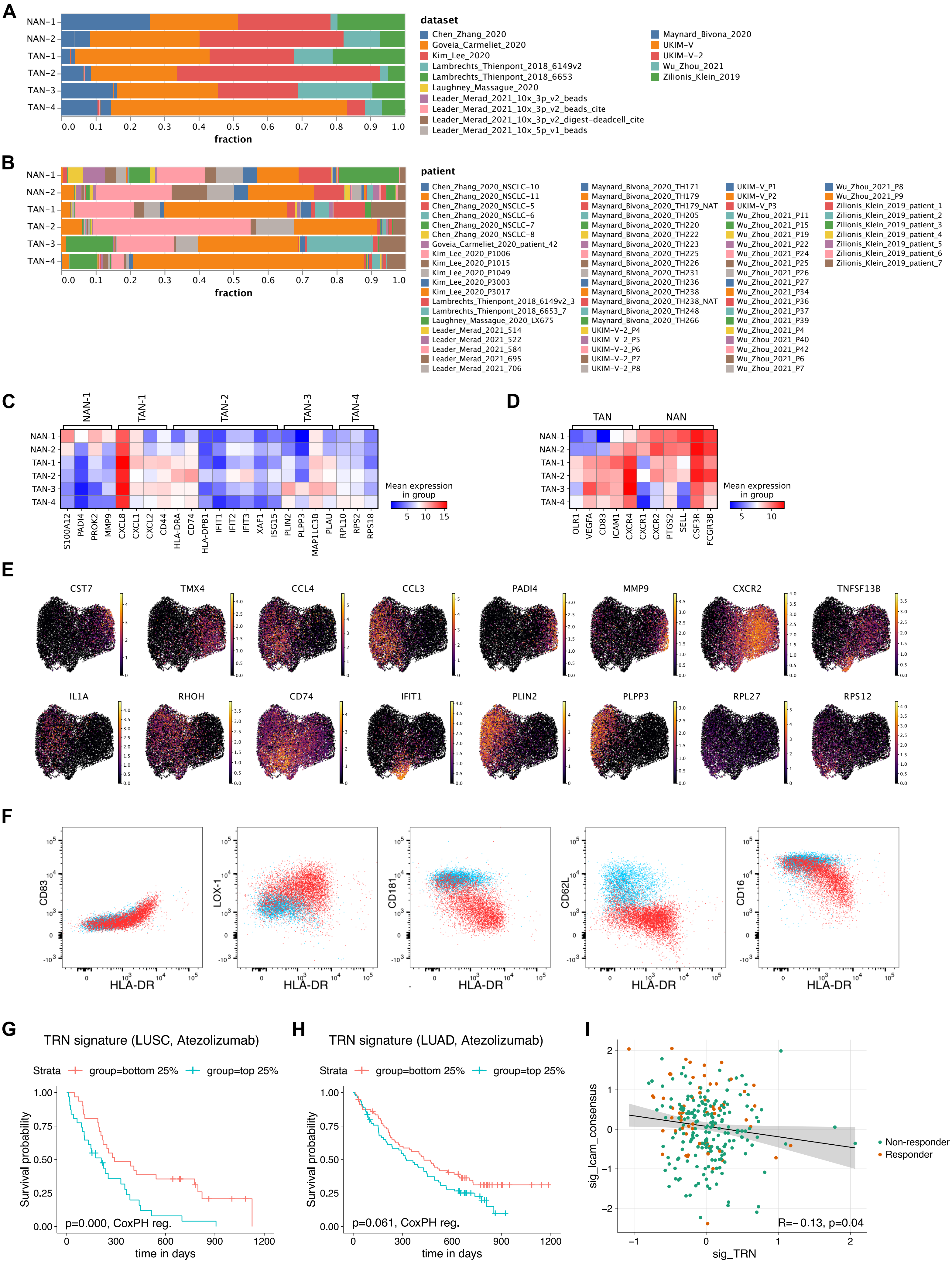

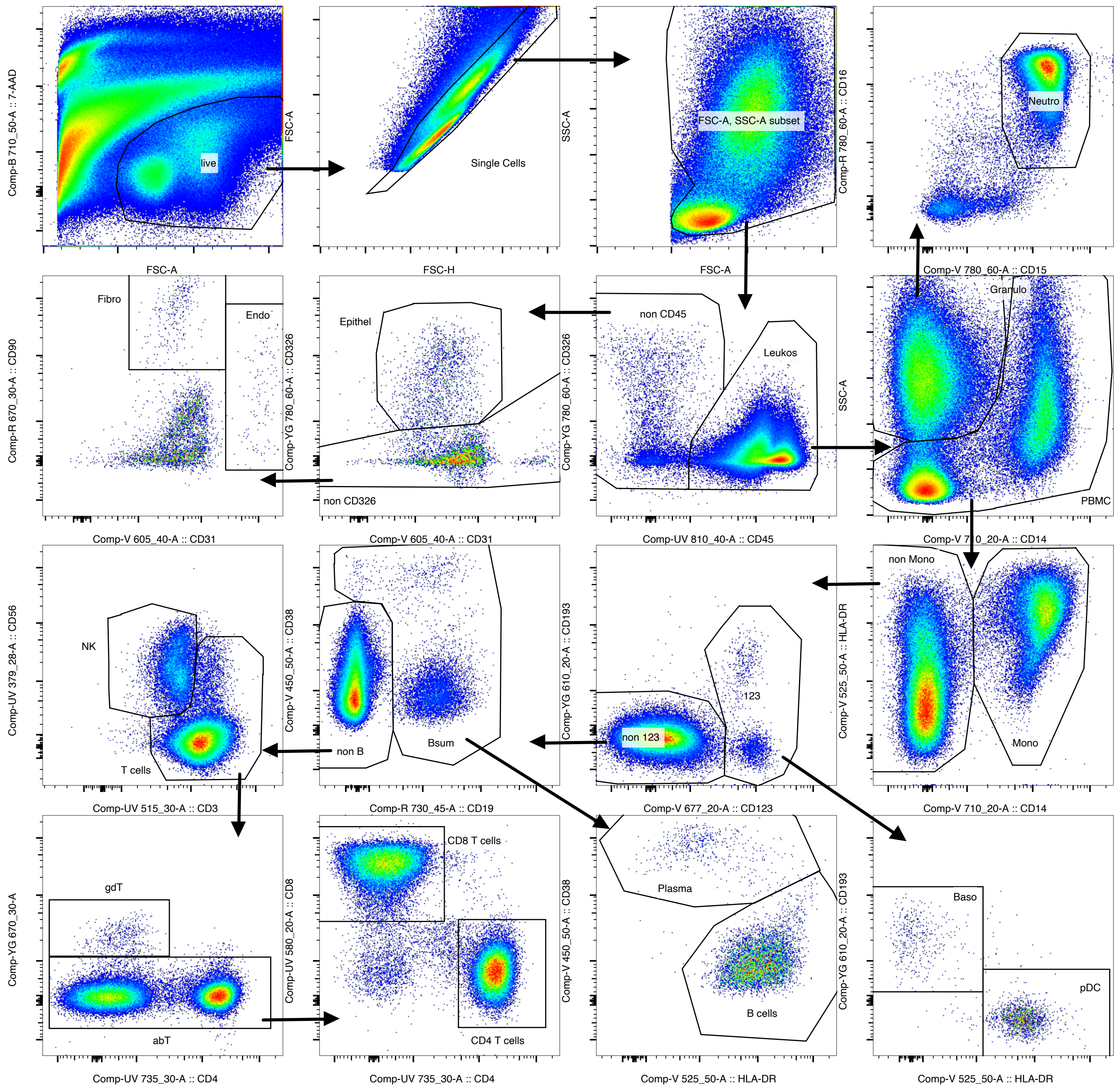
